# Bistability in oxidative stress response determines the migration behavior of phytoplankton in turbulence

**DOI:** 10.1101/2020.04.28.064980

**Authors:** Francesco Carrara, Anupam Sengupta, Lars Behrendt, Assaf Vardi, Roman Stocker

## Abstract

Turbulence is an important determinant of phytoplankton physiology, often leading to cell stress and damage. Turbulence affects phytoplankton migration, both by transporting cells and by triggering switches in migratory behavior, whereby vertically migrating cells can invert their direction of migration upon exposure to turbulent cues. However, a mechanistic link between single-cell physiology and vertical migration of phytoplankton in turbulence is currently missing. Here, by combining physiological and behavioral experiments with a mathematical model of stress accumulation and dissipation, we show that the mechanism responsible for the switch in the direction of migration in the marine raphidophyte *Heterosigma akashiwo* is the integration of reactive oxygen species (ROS) signaling generated by turbulent cues. Within timescales as short as tens of seconds, the emergent downward-migrating subpopulation exhibited a two-fold increase of ROS, an indicator of stress, 15% lower photosynthetic efficiency, and 35% lower growth rate over multiple generations compared to the upward-migrating subpopulation. The origin of the behavioral split in a bistable oxidative stress response is corroborated by the observation that exposure of cells to exogenous stressors (H_2_O_2_, UV-A radiation or high irradiance), in lieu of turbulence, caused comparable ROS accumulation and an equivalent split into the two subpopulations. By providing a mechanistic link between single-cell physiology, population-scale migration and fitness, these results contribute to our understanding of phytoplankton community composition in future ocean conditions.

**Significance Statement:** Turbulence has long been known to drive phytoplankton fitness and species succession: motile species dominate in calmer environments and non-motile species in turbulent conditions. Yet, a mechanistic understanding of the effect of turbulence on phytoplankton migratory behavior and physiology is lacking. By combining a method to generate turbulent cues, quantification of stress accumulation and physiology, and a mathematical model of stress dynamics, we show that motile phytoplankton use their mechanical stability to sense the intensity of turbulent cues and integrate these cues in time via stress signaling to trigger switches in migratory behavior. The stress-mediated warning strategy we discovered provides a paradigm for how phytoplankton cope with turbulence, thereby potentially governing which species will be successful in a changing ocean.

## Introduction

Turbulence regulates the distribution of dissolved and particulate matter in the ocean (*1*), and along with light and nutrient supply (*2*), shapes the fluid dynamical (*3*) and evolutionary niches of phytoplankton in marine ecosystems (*4*) by selecting fundamental traits such as body size and shape (*5*), life history strategies, and motility characteristics (*6*). Larger cells including diatoms benefit from turbulence through enhanced nutrient uptake (*5-7*), whereas turbulence is often detrimental for smaller motile phytoplankton, causing physiological impairment and physical damage (*8-10*).

Phytoplankton experience turbulence as fluctuations in fluid velocity gradients or ‘eddies’ (*8*), which transport and reorient cells stochastically. When coupled with motility, turbulence can create patchiness in the distribution of phytoplankton at millimeter to centimeter scales (the Kolmogorov scale) (*11, 12*), potentially impacting on populations’ ecology by modulating cells’ encounter rates and signaling. To cope with turbulence, phytoplankton can regulate lipid content, release of infochemicals, or gene expression profiles (*13*). On faster timescales, phytoplankton are able to respond to the fluid mechanical cues associated with turbulence (*14*) by regulating buoyancy (*15*) or switching migratory direction (*16*). In particular, some dinoflagellates and raphidophytes alter their direction of vertical migration when exposed to the periodic changes of orientation relative to gravity caused by turbulent eddies, leading to the emergence of a downward-migrating subpopulation among cells originally migrating upward (*16*). Vertical migration is a hallmark of many phytoplankton species (*17*), serving to harness light by day and nutrients at depth by night (*18*). The physiological mechanisms mediating the nexus between turbulence and vertical migration are thus key to understanding how the hydrodynamic environment shapes phytoplankton dynamics, in today’s oceans as well as in future, altered turbulence regimes induced by climatic changes (*19*). Yet, a fundamental understanding of the impact of turbulence on phytoplankton migratory behavior, physiology and fitness, is hitherto lacking.

Here, using a combination of millifluidics-based visualization, quantification of stress accumulation, photophysiology and growth dynamics, and mathematical modelling, we report that the emergent migratory behavior of the marine raphidophyte *Heterosigma akashiwo* exposed to turbulent cues is determined by the interplay between the Kolmogorov timescale set by the intensity of turbulence and two fundamental cellular timescales: the timescale set by a cell’s mechanical stability to overturning and the timescale of the dissipation of intracellular stress. We used time-lapse imaging to track the migration of individual cells in a small (12 mm × 4 mm × 1.6 mm) rotating chamber (*16*) that can be rotated around a horizontal axis by a computer-controlled motor with any user-defined time series of the rotation angle. We performed experiments for different rotation time series, as a model system to determine the effect of the magnitude and intermittency of small-scale turbulent eddies (*20*). Ocean turbulence is often intermittent or patchy, and its magnitude highly variable, with turbulent kinetic energy values ranging from *ε* = 10^−10^ to 10^−5^ W kg^−1^ (*5, 8*), which correspond to Kolmogorov timescales *τ*_K_ = 100 to 0.3 s. Our experimental system models intermittent turbulence as a sequence of reorientations of the chamber of magnitude π, each taking a time *τ_R_*, separated in time by a resting time *τ*_W_ during which the chamber is kept still (Fig. 1A,B). We hypothesized that the interplay between the rotation rate, Ω = π/*τ*_R_, and the time available for recovery, *τ*_W_, would regulate the emergence of the downward-migrating subpopulation from an initially upward-migrating population.

**Fig. 1.**
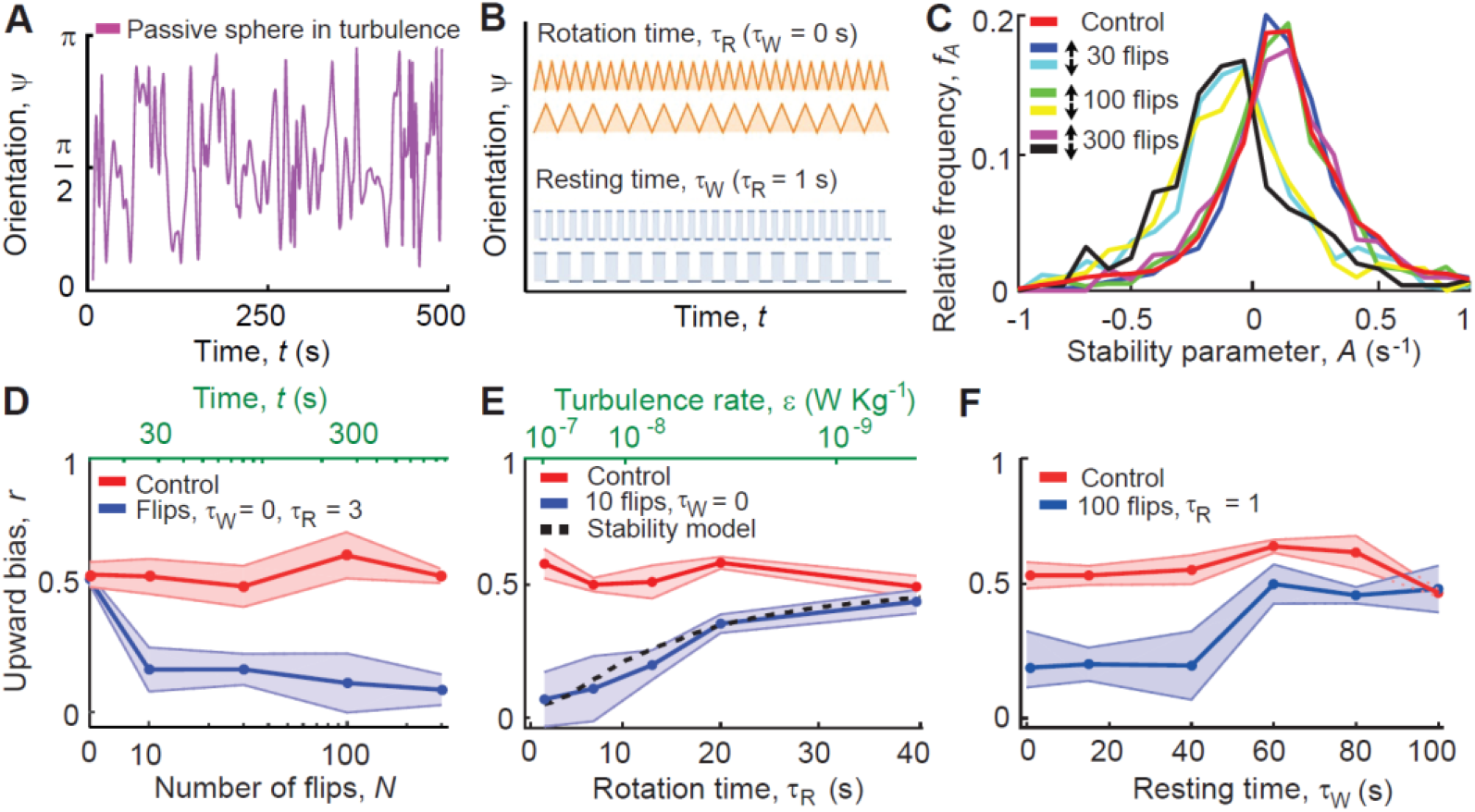
Rotation rate and resting time between reorientations relative to gravity determine the migratory response of Heterosigma akashiwo to turbulent cues. (**A**) Time series of the orientation, ψ(t), of a passive sphere relative to the vertical in a three-dimensional isotropic turbulent flow, obtained from a direct numerical simulation. The signal reveals the characteristic effects of the microscale turbulent eddies, that is, periods of time where the sphere abruptly changes its orientation by up to an angle π (modeled in this work by a rotation time), alternating with regions in which the orientation is more constant over time (modeled in this work by a resting time). (**B**) Experiments are based on a simplified characterization of intermittent turbulence, in terms of two parameters: the rotation time, τ_R_, over which the experimental chamber completes a reorientation of amplitude π (one ‘flip’ at a rate Ω = π/τ_R_), and the resting time, τ_W_, during which the chamber is held still between reorientations. Orange curves show the case with no intermittency (τ_W_ = 0 s), blue curves show two cases of short and long resting times τ_W_, respectively. (**C**) Relative distribution of the cells’ mechanical stability, expressed as the stability parameter A. The red curve corresponds to a population of cells before flipping. Other colors correspond to cells from the top (↑) and bottom (↓) subpopulations after N = 30, 100 and 300 flips (τ_R_ = 3 s; τ_W_ = 15 s). (**D**) The upward bias index, r (Methods, SI) as a function of the number of flips, N, decreases from 0.52 to 0.17 over only 30 s of flipping (τ_R_ = 3 s, time elapsed t = N τ_R_). (**E**) The upward bias as a function of the rotation time, τ_R_, for a constant resting time, τ_W_ = 0 s (blue curve). Faster reorientations (smaller τ_R_), which correspond to stronger turbulence, ε, cause a large population split, when evaluated over the same number (10) of flips. Our model of cell stability (dashed line, Methods) correctly predicts the upward bias, i.e., the fraction of downward-migrating cells that emerge for each treatment. (**F**) The upward bias as a function of the resting time, τ_W_, for a constant rotation time, τ_R_ = 1 s (blue curve). Shorter resting times (smaller τ_W_) induced a large population split, when evaluated over the same number (100) of flips. In panels D-F, circles and shaded regions denote mean ± s.d. of four replicate experiments, and corresponding controls (measured over the same time period, but without flipping) are shown in red.

## Results

The change in migratory behavior occurred within the first 10 overturning events (Fig. 1D), corresponding to only tens of seconds at the highest rotation rate used (Ω = 1 rad s^−1^, equivalent to a turbulence intensity *ε* = 10^−7^ W kg^−1^; Methods). Exposure to additional reorientations had no further effect on migration, as shown by the stable value of the upward bias index *r* over 10–300 reorientations (ANOVA, *F*_3,13_ = 0.26, *p* = 0.85). The upward bias index, *r* = (*f*_↑_ − *f*_↓_)/(*f*_↑_ + *f*_↓_), measures the relative proportion of up-swimming (*f*_↑_) and down-swimming *f*_↓_) cells (Methods). Varying the rotation rate Ω (0.08 rad s^−1^ ≤ Ω ≤ 1 rad s^−1^), for a fixed number of 10 reorientations with no resting time (*τ*_W_ = 0 s), changed the proportion of downward-migrating cells (Fig. 1E). At the fastest rotation rate tested (Ω = 1 rad s^−Γ^), the highest concentration of downward-migrating cells was observed (*r* = 0.17 ± 0.11), while at the slowest rotation rate tested (Ω = 0.08 rad s^−1^), the upward biased index (*r* = 0.44 ± 0.05) was not different from the non-rotating control experiment (*r* = 0.49 ± 0.05; *t_8_* = 0.93, *p* = 0.38) (Fig. 1E). These results show that the stronger disturbances associated with faster reorientations triggered a stronger response and more cells changed their direction of migration relative to gravity.

The initial stability of a cell regulates how the cell is affected by reorientations. The mechanical stability of a cell allows the cell to maintain its orientation with respect to gravity and is measured by the stability parameter *A* = (2*B*)^−1^, where *B* is the characteristic time for the cell to rotate back to its vertical equilibrium orientation once perturbed from it. An analysis of the stability of a cell swimming in an eddy with rotation rate Ω predicts that the cell swims at an angle *θ*_eq_ = arcsin(Ω*A*^−1^) relative to gravity if |Ω|*A*^−1^ < 1, or tumbles in a periodic orbit if |Ω|*A*^−1^ > 1 (*21*) (Methods). From this analysis we can predict the fraction of cells that switch their direction of migration from upward to downward as a function of the rotation rate Ω and the initial distribution of mechanical stabilities within a population (Fig. 1E). The latter we measured experimentally for a monoclonal population of *H. akashiwo* (CCMP452) at the single-cell level (Fig. 1C, Methods), yielding a distribution of the stability parameter characterized by high variability (*A_452_* = 0.09 ± 0.21 s^−1^). From the distribution of the cells’ initial stability parameter, our stability model correctly predicts the fraction of downward-migrating cells that emerge for each reorientation treatment (Methods, Fig. 1E). The cell’s mechanical stability thus effectively imposes a high-pass filter on the turbulence signal, whereby only reorientations on timescales *τ*_R_ shorter than the stability timescale *B* cause an upward-migrating cell to tumble (Figs. S1B, S2A) and can thus trigger the emergence of downward migration. To further test this conclusion, we performed experiments on a second *H. akashiwo* strain (CCMP3374; Fig. S1C), which has higher mean stability than CCMP452 (*A_3374_* = 0.23 s^−1^; Fig. S1A). We observed that CCMP3374 cells shift to downward swimming at higher rotation rates compared to CCMP452 cells (Fig. S1C), supporting our prediction that downward migration emerges when cells become destabilized, *i.e*., when |Ω|*A*^−1^ > 1 (Figs. S1B, S2A).

The migratory behavior was further affected by the resting time *τ*_W_, a measure of the signal’s intermittency (Fig. 1B). This was revealed by experiments with fast reorientations (Ω = 3.14 rad s^−1^), which induce population split in the absence of resting time (Fig. 1E; Methods). When the resting time was increased from *τ*_W_ = 0 s to 100 s, we found the population split to occur for values of *τ*_W_ below a threshold of 40 s (Fig. 1F), a value in line with the typical interarrival time between reorientations experienced by CCMP452 cells in strong turbulence (Fig. S2C). A threshold response is characteristic of stress responses in eukaryotes (*22*), including dinoflagellates (*23*) and diatoms (*24*), and led us to hypothesize that a progressive intracellular accumulation of oxidative compounds resulting from the reorientations is the physiological mechanism for the change in migration direction.

To test this hypothesis, we performed experiments with cells stained using a marker (CM-H_2_DCFDA) that forms a fluorescent compound in the presence of reactive oxygen species (ROS), signaling molecules that mediate the perception of diverse environmental stress conditions (*25*). Intracellular ROS accumulation was quantified by flow cytometry (Fig. S3, Methods). For these experiments, we used continuous rotation on a roller device (*τ*_W_ = 0 s, Methods) with a sample volume (2 ml) larger than the millifluidic chamber (75 μl) and thus more suitable for analysis by flow cytometry. Vertical migration was found to be independent of whether rotation was continuously in one direction (i.e., rolling) or alternating between clockwise and counterclockwise (i.e., flipping) (Fig. S4). After just 1 min of rolling, downward-migrating cells were found to have accumulated two-fold more ROS compared to upward-migrating cells (Fig. 2A). This observation indicates that a bistability in oxidative stress response underpins the split in migratory behavior of phytoplankton cells experiencing turbulent cues.

**Fig. 2.**
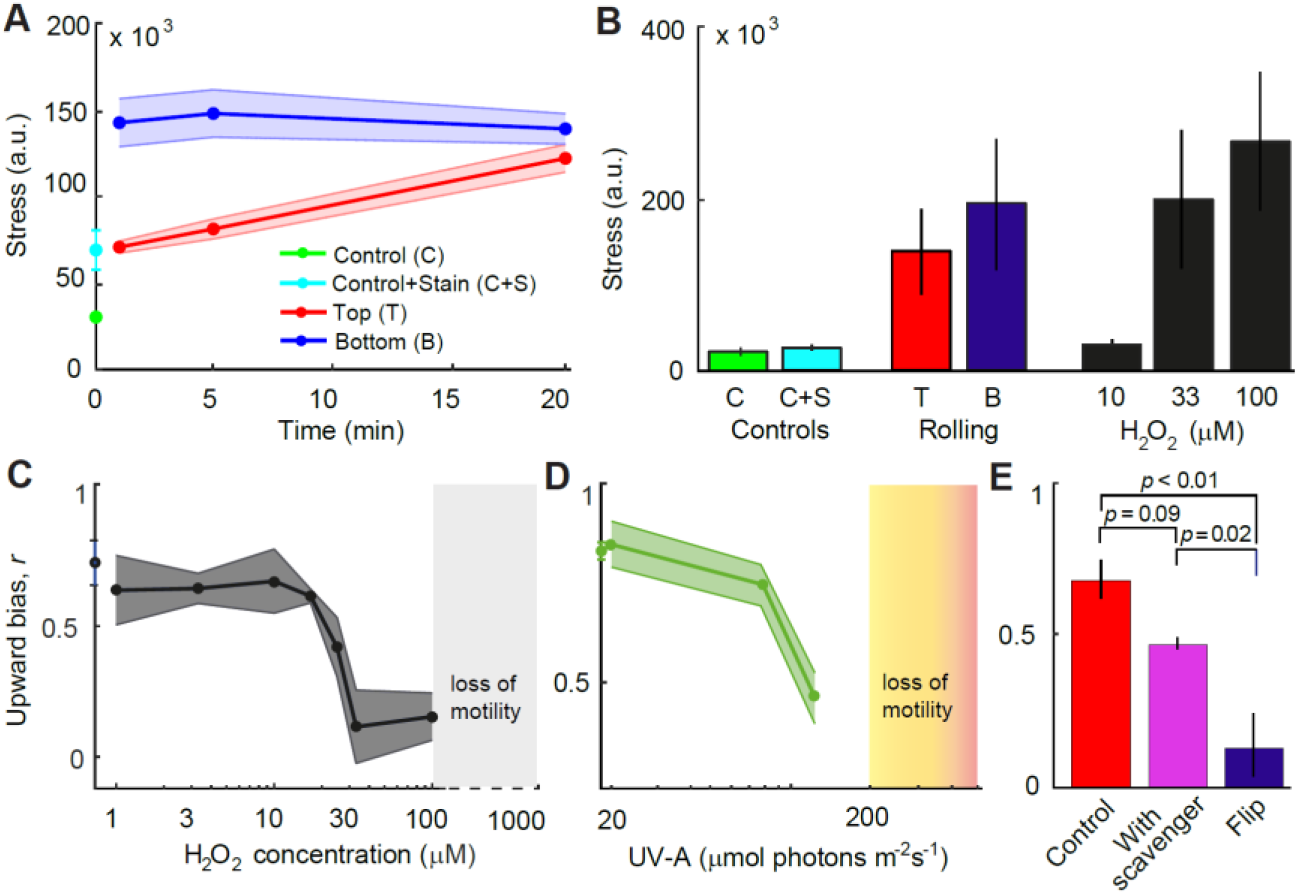
Bistability in oxidative stress mediates vertical migration of H. akashiwo. (**A**) Oxidative stress level, caused by intracellular ROS accumulation, as a function of the time exposed to rolling (Ω = 1 rad s^−1^). Curves show the increase in oxidative stress levels for the top (red, ‘T’) and the bottom (blue, ‘B’) subpopulations. Also shown are the baseline fluorescence signal of untreated control cells (green, ‘C’) and of control cells treated with the fluorescent stain CM-H_2_DCFDA, a general oxidative stress indicator (cyan, ‘C+S’). Stress levels were computed from flow cytometric measurements (mean ± s.d. of three replicates). (**B**) Oxidative stress level caused by exposure to different concentrations of exogenous H_2_O_2_ (black bar) or to 20 min of rolling (red and blue bars, denoting the top and bottom subpopulations, respectively). Controls (same as in panel A) are shown in green and cyan. (**C**) Upward bias as a function of different concentrations of hydrogen peroxide (H_2_O_2_) exogenously added to the medium 30 min before measurements. H_2_O_2_ concentrations above 15 μM elicit the population split in migration direction. Above 100 μMH_2_O_2_ cells lost motility (Fig. S6B). (**D**) Upward bias as a function of different intensities of UV-A light (emission peak = 395 nm) applied for 30 min. Photon flux densities above 80 μmol photons m^−2^ s^−1^ elicit the population split in migration direction. Above 200 μmol photons m^−2^ s^−1^ cells lost motility (Fig. S6C). (**E**) Upward bias for cells kept in still conditions (red bar, control), for cells that were flipped (blue bar, 100 flips, Ω = 1 rad s^−1^, τ_W_ = 0 s), and for cells that were flipped after having culturing in the presence of a scavenger of reactive oxygen species (100 μM potassium iodide, magenta). Populations differed significantly in upward bias (one-way ANOVA, F_2,7_ = 19.8, p = 0.001). Brackets show p-values from post-hoc Tukey’s honest significant difference (HSD) tests (Table S1). For all panels, data shown correspond to mean ± s.d. of three replicates.

To further support the finding that ROS affects migration behavior, we observed the migration of cells exposed to different exogenous stressors known to cause ROS accumulation. In a first set of experiments, we added hydrogen peroxide (H_2_O_2_) to the medium. H_2_O_2_ diffuses across the cell membrane, mimicking the physiological intracellular accumulation of ROS caused by the reorientations. Exposure to exogenous H_2_O_2_ induced the population split in migratory behavior above a threshold concentration of 15 μM H_2_O_2_ (Fig. 2C), with a threshold-like behavioral response akin to that caused by fast reorientations (Fig. 1F). The endogenous ROS levels observed upon rolling were similar to those observed upon exogenous treatment with 33 μM of H_2_O_2_ (Fig. 2B). Most notably, the ROS levels of cells increased sharply upon increasing the concentration of exogenous H_2_O_2_ from 10 to 33 μM (Fig. 2B), a concentration range that matches with the H_2_O_2_ concentration (15 μM) causing the population split (Fig. 2C). In a second set of experiments, we exposed cells to light for 30 min at intensities known to lead to ROS accumulation (*26*) and characteristic of ocean surface waters (*27*). We observed the emergence of a downward-migrating subpopulation for cells exposed to near-UV-A light (380–400 nm) at intensities greater than 80 μmol photons m^−2^ s^−1^ or to full-spectrum light (320–800 nm) at intensities greater than 650 μmol photons m^−2^ s^−1^ (Figs. 2D, S5A). Finally, we repeated the overturning experiments for cells pre-treated with the ROS scavenger potassium iodide (Methods) at an exogenous concentration of 100 μM (Fig. 2E, Table S1). No emergence of a downward-migrating subpopulation was observed in this case. Taken together, these experiments demonstrate the link between intracellular stress accumulation mediated by ROS and the behavioral switch in migration direction.

To predict the emergence of the behavioral switch in the migratory response, we devised a mathematical model of stress dynamics in cells exposed to turbulence. In the model, the cell’s mechanical stability prevents overturning by the weaker eddies (Figs. 1E, S1B, S2A), thus creating resting times between periods during which the cell is overturned (Figs. 3A, S2B,C). Accordingly, a cell accumulates ROS whenever it is tumbled by an eddy, and dissipates stress by means of its intracellular antioxidant capacity (*28*), with a characteristic dissipation timescale *τ*_S_ (see Methods, Eqs. 3 and 4). We quantified stress dissipation dynamics and the timescale *τ*S experimentally by observing the reduction of ROS over time for cells after exposure to continuous rolling for 5 min (Ω = 1 rad s^−1^). These experiments showed that stress decays exponentially over time with a timescale *τ*_S_ = 87 ± 32 s (Fig. 3B). Using this model of stress accumulation-dissipation dynamics, we predicted the time series of stress accumulation (Fig. 3C) for individual cells exposed to rapid reorientations (Ω = 3.14 rad s^−1^), for the same range of resting times τ_W_ studied experimentally (Fig. 1F). This allowed us to compute the maximum stress accumulated by cells during reorientations as a function of the resting time, and to compare this with the experimentally measured ROS concentrations at which downward-migration emerged (Fig. 2A). We find that the theoretical predictions for the maximum resting time for the emergence of downward-migrating cells (τ_W_ = 48 s, Fig. 3D) quantitatively match the values of τ_W_ observed experimentally (τ_W_ = 40 s, Fig. 1F). The model further reveals a general criterion for the emergence of downward-migration: when τ_S_/τ_W_ >> 1 (Eq. S7 in SI), the ROS scavenging machinery of a cell is too slow in dissipating stress relative to rate at which stress accumulates owing to the interarrival time of reorientations induced by turbulence, and the accumulated stress induces the switch in migratory behavior.

**Fig. 3.**
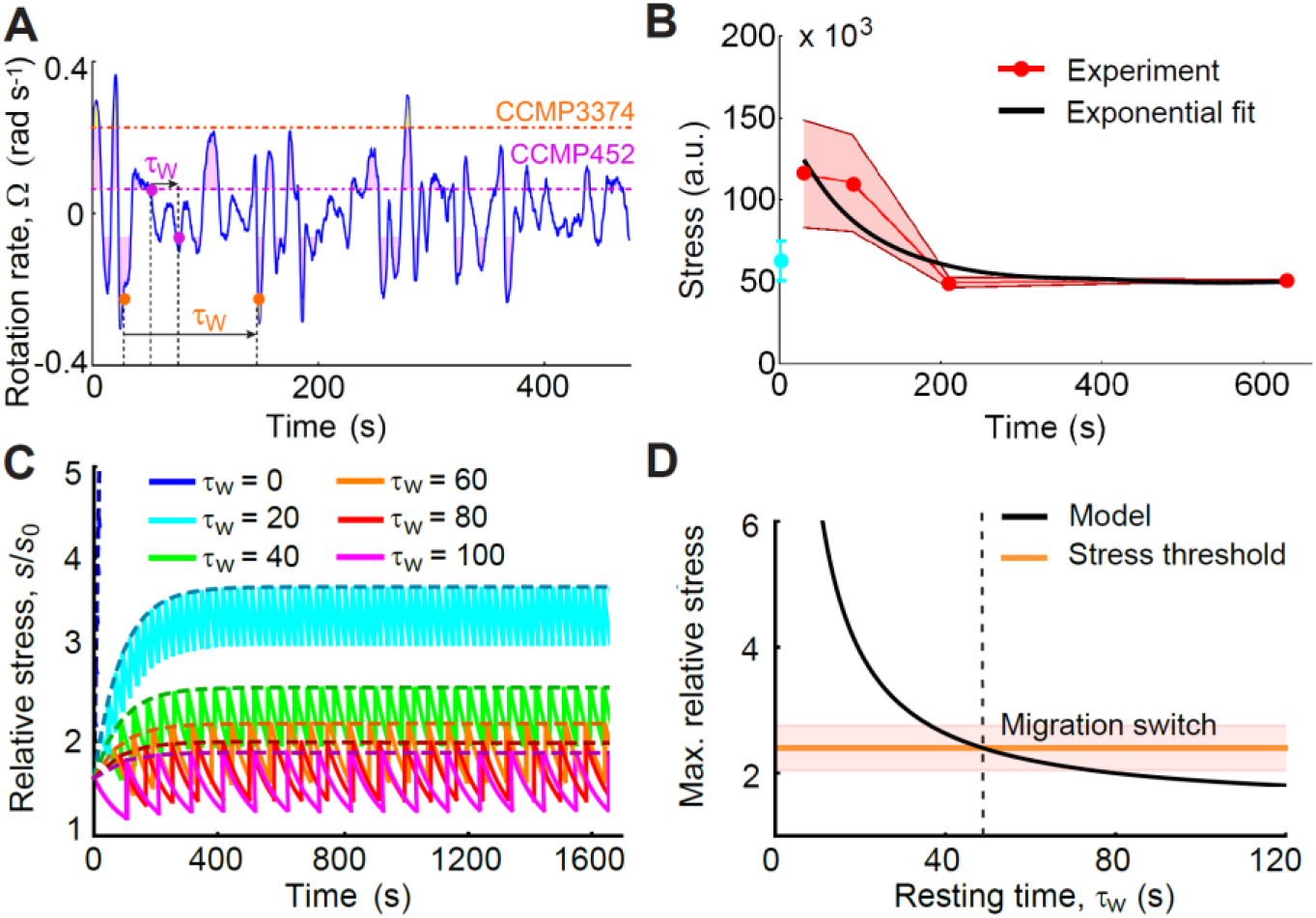
The dynamics of ROS accumulation–dissipation regulate migratory behavior of H. akashiwo. (**A**) Time series of the rotation rate relative to gravity, Ω, of a passive cell in a 3D isotropic turbulent flow, obtained from a direct numerical simulation. Dashed lines represent the value of the stability parameter, A, for CCMP452 (magenta) and CCMP3374 (orange). Shaded regions represent time windows over which a cell will be tumbled by turbulence (i.e., |Ω|A^−1^ > 1, Fig. S2A), corresponding in our experiments to imposed reorientations. During times when the rotation rate is within the two dashed lines (i.e., |Ω|A^−1^< 1), cells are not tumbled but will achieve an equilibrium swimming orientation (Fig. S1B), corresponding in our experiment to a resting time, τ_W_, between reorientations. The cell’s mechanical stability thus imposes a high-pass filter on the turbulent signal, with the higher mechanical stability of CCMP3374 resulting in longer resting times (Fig. S2B,C). (**B**) The stress dissipation dynamics, measured in still conditions (red, mean ± s.d. of four replicates) for a population previously exposed to 5 min of rolling (Ω = 1 rad s^−1^), is characterized by an exponential decay, with timescale τ_S_ = 87 ± 32 s (black). (**C**) Time series of stress (Eqs. 3–4, Methods) predicted by the mathematical model for the same range of resting times investigated experimentally (Fig. 1F). Cells rapidly accumulate stress after being reoriented (Ω = 3.14 rad s^−1^) and dissipate it with timescale τ_S_. The dashed curves represent the upper envelope of the stress signal (Eq. S5 in SI). Stress values have been normalized by the baseline stress level, s_0_, for a population under still conditions. (**D**) Predicted maximum relative stress after flipping (N = 100, Ω = 3.14 rad s^−1^), as a function of the resting time τ_W_ (black; Eq. S6 in SI). The orange line at h = 2.4 ± 0.4 corresponds to the threshold value of the stress measured experimentally for the bottom subpopulation after 1 min of rolling relative to the stained control population (Fig. 2A; mean ± s.d. of three replicates). The black line intersects with the orange line at τ_W_ = 48 s (vertical dashed line): for smaller resting times the model predicts a migration switch, corresponding to the threshold in τ_W_ for which the population split occurs in experiments (Fig. 1F).

The accumulation of ROS in response to turbulent cues directly affected cell physiology, for up to multiple cell divisions after cessation of the cue. Single-cell photophysiological measurements using pulse-amplitude modulated chlorophyll fluorometry (PAM; see Methods) showed that the downward-migrating cells emerging after 5 min of continuous rolling (Ω = 1 rad s^−1^) had 15% lower photosystem (PS) II photosynthetic quantum yields (*F_v_/F_m_*) compared to upward-migrating cells (Fig. 4A, Table S2). This reduction may stem directly from endogenous ROS, which can reduce photosynthetic quantum yields (*29*) via the general suppression of PSII D1 protein synthesis and repair (*30, 31*), activation of non-photochemical pathways, or photoinactivation of PSII reaction centers (*32*). This reduction in photosynthetic performance in *H. akashiwo* is acute when compared to the typically <10% reductions caused by high light exposure in diatoms (*26, 29*). We further found evidence for longer-term physiological damage after 5 min of continuous rolling corresponding to strong turbulence (Ω = 1 rad s^−1^), with the downward-migrating subpopulation exhibiting a 35% lower growth rate over 4 days than the upward-migrating subpopulation (Fig. 4B, Methods). This growth reduction over multiple generations (approximately four) indicates that the reorientation-induced ROS accumulation has systemic consequences for *H. akashiwo* and suggests the potential presence of a transgenerational stress memory, akin to epigenetic effects observed in plants (*33*). This result is in contrast with stress propagation in *E. coli* and yeast (*22*), where mother cells retain the oxidized aggregated protein, leaving daughter cells cleared of damaged proteins (*28*).

**Fig. 4.**
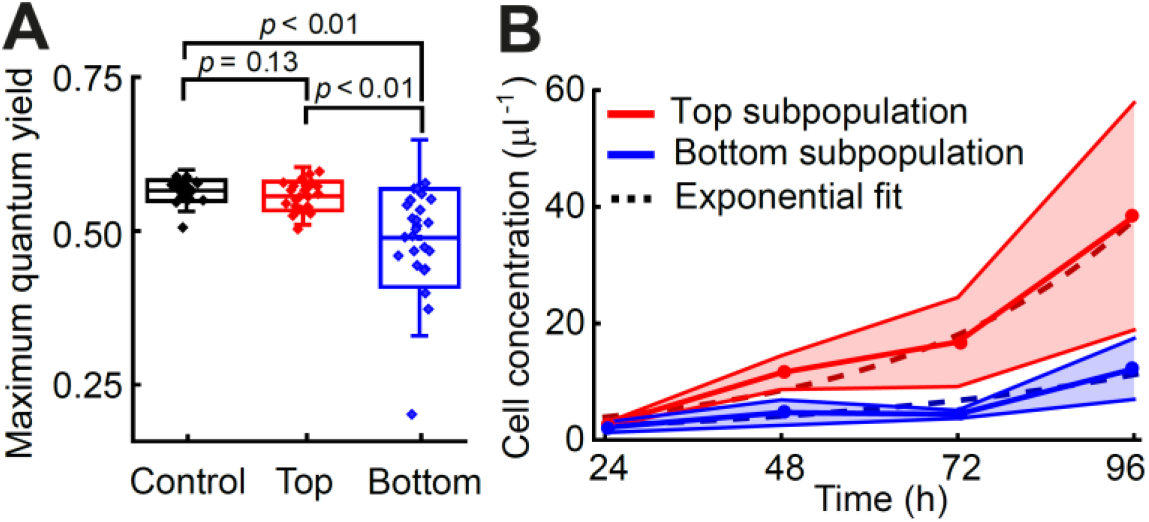
The oxidative stress induced by turbulent cues reduces photosynthetic efficiency and growth rate of H. akashiwo. (**A**) Maximum photosynthetic quantum yield, F_v_/F_m_, of H. akashiwo cells before and after turbulence-induced population splits. A pulse-amplitude modulated (PAM) chlorophyll fluorometer was used to assess the maximum quantum yield of photosystem II on single cells collected from the top (n = 26) and bottom (n = 25) of the chamber after exposure to rolling (Ω = 1 rad s^−1^) for 5 min, and from cells not exposed to rolling (control, n = 25). All cells were dark-adapted for 15 min before measurements. Boxes show ± 1 s.d., whiskers ± 2 s.d., and the central line indicates the mean. Control and flipped populations differed significantly (one-way ANOVA, F_2,74_ = 18.8, p < 0.001). Brackets show p-values from post-hoc Tukey’s honest significant difference (HSD) tests (Table S2). (**B**) Increase in cell concentration with time for the subpopulations extracted from the top (red) and bottom (blue) of the chamber after exposure to rolling (Ω = 1 rad s^−1^) for 5 min (mean ± s.d. of three replicates). Cells were regrown from the same initial density. The intrinsic growth rate, g, was quantified for each subpopulation by fitting an exponential function (dashed curves; g_↑_ = 0.74 ± 0.02 day^−1^, g_↓_ = 0.47 ± 0.03 day^−1^).

## 3 Discussion

The evidence we have presented for the role of stress in vertical migration provides a new view of the ecological implications of the active response of phytoplankton to turbulence. Our results demonstrate that the overturning of cells, a fundamental yet to date unappreciated mechanical cue due to turbulence in the ocean, can trigger behavioral and physiological responses over timescales spanning tens of seconds to multiple generations. The good agreement between our model and observations suggests that motile phytoplankton use mechanical stability to sense the intensity of turbulent cues and integrate these cues in time via ROS signaling: when ROS accumulates beyond a threshold, it triggers the switch in migratory behavior. This ROS-mediated early warning strategy may be advantageous owing to the heterogeneity in mechanical stability within monoclonal populations (Fig. 1C). The reorientations used in our experiments, corresponding to moderate to strong levels of turbulence (*ε* = 10^−9^ −10^−6^ W kg^−1^, Methods) did not inhibit motility (Fig. S6A). By responding to the ROS-mediated early warning upon first encountering a region of turbulence, cells with weaker mechanical stability will avoid swimming into the ‘eye of the storm’, where they could get trapped (*34*), damaged, or lose motility (8–*10*). Heterogeneity is also seen in the antioxidant capacity, potentially the product of a tradeoff in which cells with low antioxidant capacity are more sensitive to turbulent cues via ROS signaling, but at the cost of weaker protection against other environmental cues eliciting oxidative stress.

The emergence of downward-migration upon exposure to well-known ROS inducers (H_2_O_2_, UV-A radiation, high irradiance), in the absence of turbulent cues, shows that ROS accumulation is the cause for the migratory response, but at the same time begs the question of how specificity of ROS signaling (*25, 28*) towards turbulence might be achieved in *H. akashiwo*. In fact, specificity of response may not be necessary, if avoidance is a universally appropriate response to accumulation of ROS. Elevated levels of exogenous H_2_O_2_ (100 μM), UV-A radiation (300 μmol photons m^−2^ s^−1^), and full-spectrum light (650 μmol photons m^−2^ s^−1^) negatively impacted motility (Figs. S5B, S6B,C). Exposure to excessively high levels of irradiance can cause photoinhibition (*26, 29*), and downward migration would be a relevant response in the upper layers of the ocean, where cells can experience light exposure similar to the levels used in our experiments (*27*).

Despite the short timescales we have observed in the behavioral and physiological response of cells to turbulence (tens of seconds to minutes), the physiological ramifications and acclimation of plankton to environmental signals in the ocean can be long-term (*4*). A reduction in the growth rate of the emergent downward-migrating subpopulation indicates that even brief exposure to strong turbulence, in the order of hundreds of seconds, can induce lasting physiological changes, similarly to those induced by longer exposure to other environmental factors such as intense light (*26-29*), temperature (*35*) or nutrient availability (*2, 4, 36*). Finally, our results suggest that global warming will not only impact phytoplankton physiology and metabolism directly, but also indirectly through their response to decreased mean turbulence intensities and more energetic local storm events (*19, 36*). We propose that these changes in the physical regime of the water column will favor adaptive strategies that allow cells to cope with the stochasticity of turbulence (*37*). One example of this is the response described here for a harmful-algal-bloom-forming species, mediated by high phenotypic variability in swimming mechanics and by ROS bistability. Deepening our understanding of the physiological mechanisms underpinning these adaptive strategies, exemplified in our work through the mechanism of a ROS-mediated warning system, will thus contribute to understand responses of migrating populations and ultimately community composition in future ocean conditions.

## 4 Materials and Methods

### Cell culture and growth rate measurements

Two strains of the raphidophyte *Heterosigma akashiwo* (*38*) were examined: CCMP452 and CCMP3374 (both obtained from the National Center for Marine Algae and Microbiota, Bigelow Laboratory for Ocean Sciences, Maine USA). Cells were cultured in 50 ml sterile glass tubes under a diel light cycle (14 h light: 10 h dark; 75 μmol photons m^−2^ s^−1^) in f/2 (minus silica) medium, at 21 °C (CCMP452) or 18 °C (CCMP3374). For propagation of the cell cultures, 2 ml of the parent culture was inoculated into 25 ml of fresh medium every two weeks. Experiments with CCMP452 were carried out with monoclonal cell cultures that were grown from a single parent cell, isolated using an inoculation loop (diameter ~100 μm) developed in-house. The loop was dipped into a culture to trap a thin liquid layer and microscopy was used to select the cases with only a single CCMP452 in the layer. Each individual cell was transferred to a separate well in a 36-well plate containing fresh growth medium. Experiments were conducted between 96 h and 120 h after inoculation, which corresponds to the early exponential growth phase of the species (*16*) at room temperature (21 °C). A fixed period of the day (between 09:00 h and 15:00 h) was chosen for the experiments to rule out any possible artefact due to the diurnal migration pattern of many phytoplankton species, including *Heterosigma akashiwo* (*16, 39*). For the experiments to test the effect of scavengers on the oxidative stress-induced split, CCMP452 cells were grown with potassium iodide (KI), an H_2_O_2_ scavenger (*40, 41*). Cells in preliminary trials were grown in suspensions containing a range of KI concentrations: 1 μM, 10 μM, 100 μM, and 1 mM. All scavenging experiments reported here were performed with the 100 μM KI concentration. At this concentration, the growth rate, vertical distribution and swimming speed of CCMP452 cells matched those of the control cell culture (cells grown without KI) in the absence of turbulence treatments.

To study the long-term impact of turbulent cues on growth rates, both the top and bottom subpopulations of CCMP452 were harvested (300 μl each) from a 2 ml cell culture vial that had been exposed to turbulent cues in the form of rolling for 5 min (see **“Generation of turbulent cue: (*ii*) Rolling”** in Supplementary Information). Prior to harvesting the subpopulations, the cell culture was allowed to attain the post-turbulence stationary vertical distribution (see “**Upward bias index**” section in SI). The harvested cells were introduced in the supernatant of the initial cell culture (from which the 2 ml suspension had been taken) in a 1:12 ratio, and allowed to grow over 96 h (Fig. 4B). Cells were counted every 24 h by flow cytometer (CytoFLEX S, Beckman Coulter), and in parallel, their motility was checked using phase contrast microscopy (Nikon Ti-E, Nikon, Japan). The cell concentration of each of the subpopulations was fitted over the 96 h period using the least squares method to obtain exponential growth curves (Wolfram Mathematica v. 11.3, Champaign, IL).

### Quantification of endogenous stress production

To quantify the accumulation of reactive oxygen species (ROS), cells exposed to turbulence or static conditions were incubated under dark conditions for 30 min in 10 μM CM-H_2_DCFDA (Ex/Em: ~492–495/517–527 nm, Thermo Fischer Scientific, diluted in f/2). CM-H_2_DCFDA is a chloromethyl derivative of H_2_DCFDA that enables the detection of low concentrations of ROS. The marker is a suitable indicator for long-term quantification of ROS as it passively diffuses into live cells, and forms a highly stable fluorescent adduct when oxidized. After the 30 min incubation period, fluorescence intensities of single cells were quantified using a flow cytometer (CytoFLEX S, Beckman Coulter), in the FITC-A channel (Ex/Em: ~488/520 nm). Single-cell oxidative stress levels are represented as (relative) fluorescence units. To obtain the stress levels we subtracted the FITC-A values for the control in the absence of CM-H_2_DCFDA staining from the FITC-A values for the stained cells. This additional step ensured the subtraction from the stress measurement of the characteristic autofluorescence of raphidophytes over the green portion of the spectrum (*42*). Fluorescence levels were obtained for the turbulence-exposed population for the top and bottom subpopulations, and for the control population (no turbulence) with and without the addition of CM-H_2_DCFDA.

### Pulse-amplitude modulated chlorophyll fluorometry (PAM) experiments

PAM was used to quantify the photosynthetic performance of cells after exposure to turbulent cues. Microscopic multicolor variable chlorophyll fluorescence imaging (IMAG-RGB; Heinz Walz GmbH, Effeltrich Germany) was used to quantify the photosynthetic activity of individual cells of CCMP452. A detailed technical description of the microscope system can be found elsewhere (*43*). For PAM measurements, cells were placed into one of the channels of a prefabricated glass-bottom microfluidic chamber with depth of 100 μm (Ibidi μ-Slide VI, Ibidi GmbH, Martinsried, Germany). Using the saturation pulse method (*32, 44*), which is based on recording fluorescence yields before and during a saturating light pulse, the maximum quantum yield of photosynthetic energy conversion in photosystem (PS) II, *F_v_/F_m_* = (*F_m_* – *F_0_*)/*F_m_* was measured after a 15-minute dark incubation. Photosynthetically active radiation (PAR, 400–700 nm) was provided by RGB LEDs, which were calibrated before each experiment using a PAR light-sensor (MC-MQS micro quantum sensor, Walz, Effeltrich, Germany) connected to a light meter (ULM-500, Walz). All measurements were performed using coalesced RGB LEDs (‘white light’).

### Gravitactic cells in a fluid under solid body rotation

In Stokes flow regime (Reynolds numbers < 1), the direction in which a gravitactic cell swims is at any instant determined by the balance of viscous and gravitational torques on the cell. For gravitactic cells characterized by a stability parameter *A* swimming at low Reynolds numbers in a fluid under solid body rotation in the vertical plane at a constant rotation rate Ω = π/*τ*_R_ (rad s^−1^), which here exemplifies the characteristic reorientation rate 1/*τ*_K_ by Kolmogorov-scale turbulent eddies in the ocean, the equation of motion in the laboratory frame of reference reads

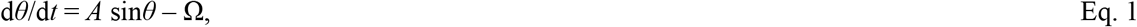

where *θ* measures the cell orientation to the vertical (Figs. S1B, S2A), and we have used the relation between the vorticity and the rotation rate for a fluid in a continuous (clockwise) solid body rotation in the vertical plane performed by the flipping chamber (and the rolling device), with the strain rate set to *E* = 0 (*21*). A gravitactic cell may therefore swim at a non-zero angle *θ*_eq_ = arcsin(*A*^−1^Ω) relative to the vertical if *A*^−1^ |Ω| < 1, with 0 < *θ*_eq_ < π/2, or it may tumble if Ω_c_ is sufficiently large (*A*^−1^|Ω_c_| > 1) and thus perform a periodic orbit with period

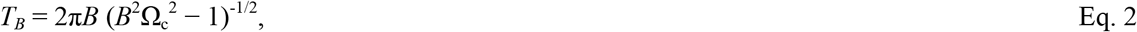

where *B* = (2*A*)^−1^ is the stability timescale (*45*). The solutions for Eq. 1 are portrayed in Fig. S1B for Ω = 0.2 rad s^−1^ and two different stability parameters corresponding to the two strains CCMP452 (low stability, where cells tumble) and CCMP3374 (high stability, where cells swim at an equilibrium angle *θ*_eq_). By applying the condition for tumbling *A*^−1^|Ω_c_| > 1 for the distribution of the stability parameter *f_A_*, measured experimentally (Fig. 1D), the fraction of cells migrating upward *f_↑_* and downward *f_↓_* can be extracted (see SI). We can then derive the upward bias *r* as a function of the rotation time *τ*_R_ of the chamber. The result of this stability analysis is plotted in Fig. 1E (black dashed line).

### Stress dynamics of gravitactic cells under turbulent cues

Above some critical value of rotation rate, Ω_c_ at which *A*^−1^ |Ω_c_|> 1, the cell tumbles by fluid shear in the vertical plane and perform periodic orbits because of a low mechanical stability (Fig. S1B). An effect of rotation is that the gravitational acceleration g appears to be rotating in the rotating frame of reference. During a tumbling event, intracellular stress is generated in the cell under the form of a nearly instantaneous release (*i.e*., a spike) of ROS whenever the cell is being reoriented relatively to gravity, that is, when the cell experiences an impulsive force of typical magnitude *F*_g_ ~ 1 pN (see SI). In our model, the ROS spike specifically occurs at times *t_i_* whenever the cell swims in a direction *θ* = π/2, that is in the direction perpendicular to the gravity vector. This particular choice of swimming direction is arbitrary, and we could choose any value between π/2 < *θ* < π without changing our results. The intracellular scavenging machinery of the cells dissipates the accumulated stress *s* with a characteristic timescale, *τ*_S_ (measured experimentally, Fig. 3B; see SI). In principle, the characteristic recovery timescale *τ*_S_ depends on the cellular antioxidant capacity: compare the response to flipping for two populations with and without the additional scavenging supplement of KI, Fig. 2E). However, all the turbulence experiments were performed with cells grown under the same conditions at a fixed period of the day to avoid diurnal fluctuations in population physiology (*46*), and we do not expect the antioxidant capacity to vary over the relatively short experimental timescales (<20 min rolling, Fig. 2A). We therefore assumed the stress dissipation timescale *τ*_S_ as a constant parameter in our model. The resulting intracellular stress accumulation–dissipation dynamics are captured by the following differential equation

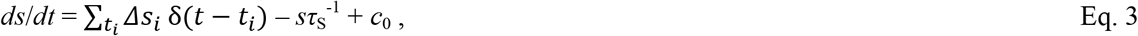

where the Dirac delta function δ(*t − t_i_*) records the stress spikes Δ*S_i_* (assumed to all have the same value Δ*s*) occurring at times *t*_i_ for a given swimming trajectory, and *c*_0_ is the baseline stress rate. Eq. 3 can be solved by performing the Laplace transform, which gives the stress level as a function of time

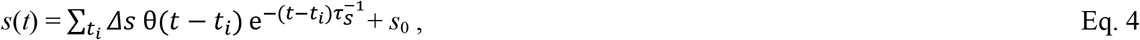

where θ(*t − t_i_*) is the Heaviside function, and *s*_0_ = *c*_0_ *τ*_S_ is the baseline stress level before the fluid rotation. The population stress levels were measured with a flow cytometer for controls (no rotations) and after rolling (Fig. 2A). We identified the stress threshold *h*, above which a cell would switch its migratory strategy, by expressing the stress *s* after exposure to rolling as the ratio *s*/*s*_0_, where *s*_0_ is the baseline stress level. The stress threshold *h* = 2.4 ± 0.4 (mean ± s.d. of three replicates), which corresponds to the orange line in Fig. 3D, is extracted from the experiment employing the shortest exposure to rolling experimentally tested (1 min rolling, which corresponds to 10 full revolutions at Ω = 1 rad s^−1^), by taking the ratio of the stress *s* for the subpopulation at the bottom (data taken from Fig. 2A, point at 1 min from the blue line B, with a fluorescent value 142 × 10^3^ a.u.) relative to the control with the same concentration of stain *s*_0_ (cyan dot, C+S, 75 × 10^3^ a.u.) after subtraction of the characteristic autofluorescence of raphidophytes over the green portion of the spectrum (green dot, C, 28 × 10^3^ a.u.). In our model, cells switch motility for *s*(*t*)/*s*_0_ > *h*. By taking the partial sum in the summation in Eq. 4, the upper envelope of the stress signal over time experienced by the tumbling cells after *N* periodic reorientations is

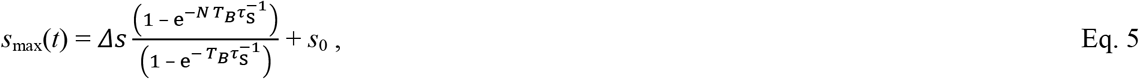

where the sequence of times *t*_i_ at which stress is generated is *S* = {*T_B_,…iT_B_,…,NT_B_*}, *T_B_* is the period of the orbit for the tumbling cells given in Eq. 2, which depends on the stability parameter *A* and on the rotation rate Ω. In the Supplementary text, we further model the stress dynamics for cells exposed to turbulent cues for the two paradigmatic cases that we employed experimentally: *i*) a continuous solid body rotation (i.e., rolling) with rotation rate Ω = π/*τ*_R_ (no resting phases, *τ*_W_ = 0), and *ii*) multiple, fast reorientations of amplitude π (i.e., flipping) occurring at a rate Ω >> *A*, alternating with resting phases captured by the timescale *τ*_W_ (Fig. 1B).

## Acknowledgements

We thank G. Boffetta and M. Cencini for sharing the direct numerical simulations data, and Russell Naisbit for help with the editing of this manuscript. This work was supported by Gordon and Betty Moore Marine Microbial Initiative Investigator Award GBMF3783 (to R.S.), and Simons Foundation Grant 542395 (to R.S.) as part of the Principles of Microbial Ecosystems Collaborative (PriME), the Israeli Science Foundation (ISF) (grant # 712233) (to A.V.), Human Frontier Science Program Cross Disciplinary Fellowship LT000993/2014-C (to A.S.) and the ATTRACT Investigator Grant A17/MS/11572821/MBRACE of the Luxembourg National Research Fund (to A.S.), and funding from the Science for life Laboratory (to L.B.).

## Author contributions

All authors designed research. A.S. performed the flipping experiments. F.C. and A.S. performed the rolling, flow cytometry, and growth experiments. L.B. performed the light and PAM experiments. A.V. designed and conceptualized the ROS signaling experiments. F.C., A.S., L.B. analyzed data. F.C. developed the mathematical model. F.C., A.S., and R.S. wrote the paper with support from all authors.

## Materials and Methods

### Generation of turbulent cues

Two different experimental modes – *flipping* and *rolling* – were employed to generate turbulent cues for the experiments reported here. The flipping mode, carried out in a flip chamber, was optimal for visualizing swimming cells and their distribution in the vertical plane, whereas the rolling mode was well suited for experiments that required larger cell culture volumes, such as those requiring subsequent fluorescent analyses using flow cytometry. Both experimental modes produced equivalent turbulent cues and yielded no difference in the turbulence-induced phytoplankton response (Fig. S4). While the rolling mode consisted of a continuous reorientation of the confined suspension in one direction, in the flipping mode, the direction alternated between clockwise and counterclockwise. The turbulent signal can be decomposed into triangular waves for reorientation occurring over different rotation time, *τ*_R_, and square waves for different resting times between reorientation, *τ*_W_ (Fig. 1A,B). In our experiments, for instance, periodic flipping consisting of multiple, rapid overturning of the chamber (180° in 3 s; *τ*_R_ = 3 s), each followed by 15 s at rest (*τ*_W_ = 15 s), results in a period of 18 s. This corresponds to the Kolmogorov timescale *τ*_K_ = (*v/ε*)^1/2^, where *v* is the kinematic viscosity of the fluid (seawater), and *ε* is the turbulent energy dissipation rate. The turbulent kinetic energy dissipation associated with *τ*_W_ = 15 s and *τ*_R_ = 3 s is *ε* = 3 × 10 W kg, a value typical of the ocean pycnocline, falling within the typical range of values for ocean turbulence (10^−10^−10^−5^ W kg^−1^) (*S1, S2*). Similarly, the turbulent kinetic energy dissipation corresponding to *τ*_W_ = 0 s and *τ*_R_ = 3 s is *ε* = 10^−7^ W kg^−1^. The two different generators of turbulent cues are described below.

### (i) Flipping

A millifluidic flip chamber (12 mm × 4 mm × 1.6 mm) made of transparent acrylic was used to visualize the motility and upward bias of *H. akashiwo* cells. The flip chamber was mounted to the shaft of a stepper motor that allowed for full reorientations from 0° to 360°. For the flipping experiments, the rotation of the chamber was automated using an externally programmed controller that drove the motor, with full user control over the time series of the rotation angle. For all experiments (including experiments with cells exposed to exogenous H_2_O_2_, UV-A, strong radiation, and after treatment with ROS scavenger KI), a 75 μl suspension of phytoplankton cells was gently pipetted into the chamber through one of two injection ports, which were then closed with silicone plugs. During experiments, cells in the flipping chamber were visualized using a stereoscope (Nikon SMZ1000) with a Plan APO × 1 objective (0.12 NA) and a digital CMOS camera (Photron FastCam SA3). The flipping chamber was mounted on a translation stage, the position of which could be controlled using micrometer screws along all three axes. The camera was focused on a plane perpendicular to the rotation axis and midway between the two chamber walls. The depth of focus was 750 μm, ensuring that cells were more than 400 μm (>50 cell radii) from the front and back walls of the chamber, to eliminate wall effects. Any small residual wall effects that may still have occurred would have been present for the entire duration of an experiment, and thus could not have caused the population split. Images were acquired at 60 frames per second. The suspension was uniformly illuminated using a single 627 nm LED (SuperBright LEDs, RL5-R12008, 0.1 W) mounted just outside of the flipping chamber. Neither of the two *H. akashiwo* strains tested showed any phototactic bias to wavelengths of light in the red spectrum, in agreement with literature (*S3*). All experiments were conducted under diffused room light settings, to avoid possible photo-responses. For each treatment, a control experiment was performed in which cells were observed in the flipping chamber without rotation, for the same duration as the treatment. The vertical distribution of cells in these control experiments was quantified at regular intervals to confirm that the upward bias of cells in the absence of overturning remained constant.

### (ii) Rolling

Turbulence experiments for subsequent flow cytometry and microscopy-PAM measurements (see below) were carried out using a programmable rolling device (Thermo Fischer Scientific Tube Roller). Cell suspensions were pipetted into cylindrical glass vials of 2 ml maximum volume, filling them to capacity to leave no empty space. The glass vials, placed horizontally on the tube roller, underwent rolling for a pre-assigned duration at a desired angular speed. The turbulence energy dissipation rates of the rolling experiments at 10 rev min^−1^ (Ω = 1 rad s^−1^) quantitatively matched those in the flipping experiments at a rotation time of *τ*_R_ = 3 s (Fig. S4). Post rolling, the cylindrical vial was left undisturbed for 30 min in the vertical position, so that the swimming cells could attain their stationary population distribution. The top and bottom subpopulations were then harvested for subsequent analyses and physiological experiments. As in the case of the flip experiments, all rolling experiments were conducted under diffused room light settings, and for each treatment, a control experiment was performed in which cells were observed in a cylindrical vial without rolling for the same duration as the treatment. The vertical distribution of cells in these control experiments was quantified at regular intervals to confirm that the upward bias of cells in the absence of rolling remained constant.

### Upward bias index

After the end of every flipping experiment, we allowed the population to reach its equilibrium distribution over the vertical by waiting 30 min. This period was chosen conservatively based on the observation that the concentration profile already stabilized after ≈5 min, and the consideration that a cell migrating upward at *v* = 88 μm s^−1^ (the mean swimming velocity, Fig. S6A) would cover the depth of the flipping chamber (4 mm) in less than 2 min. To quantify the migration behavior of the cells, we first obtained histograms of normalized cell concentration in the flipping chamber, within the region captured by the camera (4 mm × 4 mm) in the mid-chamber plane of focus. To quantify the asymmetry in cell distribution over the vertical, we computed the upward bias *r* = (*f*_↑_ − *f*_↓_)/*f*_↑_ + *f*_↓_), where *f*_↑_ and *f*_↓_ are the numbers of cells in the top 400 μm and the bottom 400 μm of the chamber, respectively (following the method developed in ref. *S4*). A symmetric distribution of cells corresponds to *r* = 0, whereas preferential upward-migrating corresponds to *r* > 0 and preferential downward-migrating to *r* < 0. After the flipping experiments, the two subpopulations of *H. akashiwo* migrating upward (↑) and downward (↓) were used in measurements of the mechanical stability (Fig. 1C). Control experiments consisted of cells held in the chamber for the entire duration of the flipping experiments without flipping.

### Cell tracking

To extract the swimming behavior of cells (swimming speed and stability), movies were recorded at 12 frames per second. For tracking, cell locations were determined by image analysis based on intensity thresholding using MATLAB (MathWorks) routines (*S5*). Cell trajectories were assembled by linking the locations of cells in subsequent frames, based on proximity and kinematic predictions from previous time steps, using automated software. Cells in the flipping chamber swam in helical patterns, characteristic of many motile phytoplankton species (*S6*). However, the helical component was averaged out using a 1-s moving average to reduce noise in the calculation of the stability parameter *A*.

### Quantification of the cell stability parameter

To determine cell stability at the population level for the *H. akashiwo* CCMP452 and CCMP3374 strains, we quantified the rotation rate *ω* of cells as a function of their orientation *θ* relative to the vertical. This is an established method for quantifying the reorientation timescale *B*, as greater stability will cause faster reorientation towards the stable orientation after a cell is perturbed. The greater the magnitude of *B*, the lower the mechanical stability, with the sign of *B* denoting the upward (*B* > 0) or downward (*B* < 0) stable swimming direction. The stability parameter is then calculated as *A* = (2*B*)^−1^ (ref. *S7*). To this end, we tracked individual cells over 15 s immediately following a single flip (which provided the perturbation from the stable orientation), and averaged their rotation rate over all cells as a function of *θ*. The resulting data for *ω*(*θ*) were fitted well by a sinusoidal function of the form *a* sin(*θ* + *κ*), with *a* the amplitude of the sinusoid, and where we imposed a phase shift *κ* equal to π for the upward-migrating cells (simultaneously fitting both *a* and *κ* showed consistent results for this approach). We determined the reorientation timescale (Fig. S1A) from the best-fit sinusoid as *B* = (2*a*)^−1^ sin *κ*. To account for heterogeneity in the stability, which would result in some cells reorienting faster than others, we also quantified the stability parameter *A* for CCMP452 cells at the single-cell level. From among the trajectories analyzed to extract the population stability parameter, we selected those that had data points with the orientation *θ* between −π/3 and π/3. This allowed us to extract the stability parameter from the trajectory in the approximately linear region of the sinusoidal dependence. Using this approach, we determined the distribution of the parameter *A* for cell populations from each of the different treatments (control, and following *N* = 30, 100, 300 flips for cells from the top and bottom subpopulations; Fig. 1C).

### Flow cytometry data analysis

*H. akashiwo* cells were identified by gating a suitable region in the 2D scatter plot using the channels APC-A (allophycocyanin, red fluorescence at 763/43 nm) vs. FSC-A (forward scatter, correlating with cell size, ref. *S8*). This gating was kept fixed for all subsequent experiments and ensured that only viable (i.e., red autofluorescing) *H. akashiwo* cells of a certain size were included in subsequent analyses (thereby excluding bacteria and other debris). Using the above gating strategy, FITC-A fluorescence was recorded across different treatments and the mean values of the different replicates extracted. The standard deviation associated with the measurements represents the variation of the mean values of the FITC-A channel across different replicates.

### Recovery after endogenous stress

To quantify the ability of *H. akashiwo* to recover from stress, four replicates were extracted from the same culture, introduced to 2 ml glass vials, and exposed simultaneously to 5 min of reorientations on the tube roller at 10 rev min^−1^ (equivalent to a fast rotation timescale of *τ*_R_ = 3 s, which is equivalent to a rotation rate, Ω = 1 rad s^−1^). After the rolling stopped, CM-H_2_DCFDA was added at a final concentration of 10 μM. The oxidative marker was introduced at four different time points, *t* = {30, 90, 210, 630} s, in the four different populations. All samples were incubated under dark conditions for 30 min and subsequently FITC-A fluorescence intensities were quantified using a flow cytometer. An exponential curve was fitted to the mean values of the replicates, weighted by their standard deviation, to obtain the recovery timescale *τ*_S_.

### Exogenous H_2_O_2_ exposure experiments

Hydrogen peroxide (H_2_O_2_), a common oxidative species, was used to study the behavioral response of *H. akashiwo* to an exogenous oxidative stressor. Hydrogen peroxide solution (30% (w/w) in H_2_O, Sigma-Aldrich), was dissolved in the CCMP452 cell suspension to achieve final concentrations spanning four orders of magnitude, from 1 μM to 1 mM. Cells were incubated in either 0 (control), 1, 3, 10, 15, 23, 33, 100 μM or 1 mM H_2_O_2_ for 15 min. The cells were then carefully transferred to the flip chamber and allowed an additional 15 min to reach their stationary swimming distribution. Using the imaging protocol described above (see “**Generation of turbulent cues: (*i*) Flipping**”), the upward bias was calculated from the equilibrium vertical distribution.

### Upward bias induced by near UV-A exposure and by strong irradiance

*H. akashiwo* cells were carefully pipetted into 24-well plates (Corning inc., Corning, NY, USA) using an enlarged (i.e., cut) pipette tip. The filling volume (1 ml) was chosen to achieve a maximum depth of 5 mm to avoid self-shading effects within the cell suspension. The 24-well plate was placed onto black masking tape (Thorlabs, Newton, NJ, USA) to avoid back scattering and irradiance was provided to a single well by a collimation lens (Thorlabs) connected to a light source via a liquid light guide. For illumination, two different light sources were used, (*i*) an LED-based Lumencor Spectra X light engine (Lumencor, Beaverton, OR, US) for targeted irradiance with UV-A (380–400 nm), and (*ii*) a mercury-based Intensilight light source (Nikon Corp., Tokyo, Japan) for full spectrum irradiance (320–800 nm, peak wavelengths at 407 nm, 429 nm, 534 nm and 575 nm). Different levels of photon quanta were achieved by the automated insertion of neutral density filters (Intensilight) or the regulation of voltage (Spectra X light engine). Full spectrum irradiance was calibrated via a cosine-corrected mini quantum sensor (MQS-B, Walz) placed at the level of the top of the culture suspension and connected to a light meter (ULM-500, Walz). In a similar fashion, near UV-A irradiance levels were quantified using a GigaHertz BTS256-EF luxmeter (Gigahertz-optics GmbH, Türkenfeld, Germany), using a spectral cut-off matching that of the light source (380–415 nm) and by exporting both photon flux densities in units of μmol m s^−1^ and unweighted UV-A irradiances. Displayed photon flux densities (20, 78 and 116 μmol m^2^ s^−1^, Fig. 2D) correspond to unweighted UV-A irradiances of 16, 61 and 85 W/m^2^ , respectively. Cells within 24-well plates were exposed for 30 minutes to near UV-A of different intensities and following exposure, cells were carefully transferred to the millifluidic flip chamber and their upward bias quantified using the methods described above. Corresponding control cells were placed in the same 24-well plates, but covered with a dark foil to avoid exposure to UV-A.

### Statistical analysis

We performed a one-way ANOVA to compare the upward bias index *r* among the still control and cells from two populations having undergone reorientations: cells grown in the presence of scavenger potassium iodide, and cells grown under the standard conditions. We made multiple comparisons using a post-hoc Tukey’s HSD test (Table S1). The same statistical analysis was conducted to compare the maximum quantum yields among the still control and cells from top and bottom subpopulations after the exposure to turbulent cues (Table S2). Experiments were performed with at least three replicates, with the number of replicates for each experiment reported in figure captions. All the replicates in our experiments were biological replicates.

## Supplementary Text

### Stress dynamics of gravitactic cells under turbulent cues

An effect of rotation is that the gravitational acceleration g appears to be rotating in the rotating frame of reference, i.e., for cells that tumble and perform periodic orbits because of a low mechanical stability. These changes in the cell orientation relative to gravity may cause the organism to behaviorally respond by switching the migratory direction from negative to positive gravitaxis, where the cue for the behavioral differentiation is not the direct effect of fluid velocity gradients (e.g., shear), but rather the eddies inducing changes in the cell orientation relative to gravity (*S4*). Above some critical value of Ω_c_ at which *A*^−1^ |Ω_c_|> 1, the cell tumbles by fluid shear in the vertical plane (Fig. S1B). For simplicity’s sake, we assume that before a tumbling event induced by flipping or rolling, the cell swims at its equilibrium position, that is, for negatively gravitactic cells such as *H. akashiwo*, the direction opposite to gravity, at which the angle *θ* = 0 relative to gravity. We can write the force-free condition along this direction of swimming (but a similar argument holds for any swimming direction) as *P* + *B* – *G* – *D*_↑_ = 0, where *P* is the propulsion force from the beating of the flagellum, *B* = *ρ_f_Vg* and *G* = *ρ*_p_*V*g are the buoyancy and gravity forces acting on the cell respectively with *V* the volume of the cell, *ρ*_f_ and *ρ*_p_ the densities of the fluid and of the cell, g is the gravity acceleration, and *D*_↑_ is the drag force directed in the opposite direction to the cell swimming velocity when the cell’s swimming is at the angle *θ* = 0 relative to gravity. After a tumbling event, for example induced by flipping or rolling, the cell’s swimming velocity is directed at an angle *θ* = π relative to gravity. At the downward-swimming orientation, the force-free condition is −*P* + *B* − *G* + *D*_↓_ = 0, where the propulsion *P* and the drag force *D*_↓_ switched sign as a consequence of the change in the swimming direction relative to gravity due to the reorientation. If we assume that the propulsion does not change as a consequence of the tumbling event, by combining the two conditions it gives the difference in the drag force experienced by the cell, *F*_g_ = *D*_↓_ − *D*_↑_ = 2 (*G* − *B*). By generalizing to any direction *θ* relative to gravity upon a tumbling event, a cell is therefore subjected to an impulsive force of magnitude *F*_g_ = (*ρ*_p_ − *ρ*_f_)g*V* (1 − cos*θ*). For *H. akashiwo*, which is characterized by an excess density (*ρ*_p_ − *ρ*_f_) ~ 50 Kg m^−3^ and an equivalent radius *R* ~7 μm (*S4, S9, S10*), the impulsive force generated by the gravity force during a tumbling event is *F*_g_ = 1.5 pN at *θ* = π. The value for *F*_g_ is considerably larger than the force exerted by fluid shear *F*_shear_ = *νρ*_f_*S ∂u*/*∂z* (ref. *S2*), where *v* = 10^−6^ m^2^ s^−1^ is the kinematic viscosity of seawater, *S* is the surface of the cell, and *∂u/∂z* is the shear experienced by the cell under turbulence. For example, at a turbulence dissipation rate *ε* = 10 W kg, cells are subjected to an average shear *∂u*/*∂z* ~ 0.1 s^−1^, at which *F*_shear_ = 0.06 pN, which is 25 smaller than *F*_g_. The two forces *F*_g_ and *F*_shear_ have equivalent magnitude for very high levels of ocean turbulence, *ε* = 7 × 10^−6^ W kg^−1^ (*S1, S2*), which are never reached in our reorientations experiments. In our model of stress dynamics, we therefore focus on the effects of the impulsive force *F*g on the ROS accumulation-dissipation dynamics, and we neglect the small contribution from the shear force. We hypothesize that during a tumbling event intracellular stress is generated in the cell under the form of a nearly instantaneous release (*i.e*., a spike) of ROS whenever the cell is being reoriented relatively to gravity, that is, when the cell experiences a force of typical magnitude *F*_g_. In our model, the ROS spike specifically occurs at times *t_i_* whenever the cell swims in a direction *θ* = π/2, that is in the direction perpendicular to the gravity vector. This particular choice of swimming direction is arbitrary, and we could choose any value between π/2 < *θ* < π without changing our results. The intracellular scavenging machinery of the cells dissipates the accumulated stress *s* with a characteristic timescale, *τ*_S_ (measured experimentally, Fig. 3B; see “**Recovery after endogenous stress**” above). By way of example, the scavenging machinery in the raphidophyte *Chattonella marina* – a harmful-algal-bloom species which exhibited a migratory response to turbulence similarly to *H. akashiwo* (*S4*) – comprises several antioxidant enzymes, including glutathione peroxidase, peroxiredoxin, catalase, and ascorbate peroxidase (*S11*). The resulting intracellular stress accumulation–dissipation dynamics are captured by the following differential equation

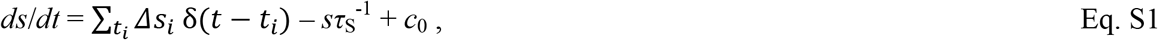

where the Dirac delta function δ(*t − t_i_*) records the stress spikes Δ*s_i_* (assumed to all have the same value Δ*s*) occurring at times *t_i_* for a given swimming trajectory, and *c*_0_ is the baseline stress rate. Eq. S1 can be solved by performing the Laplace transform, which gives the stress level as a function of time

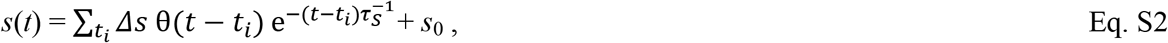

where θ(*t − t_i_*) is the Heaviside function, and *s*_0_ = *c*_0_ *τ*_S_ is the baseline stress level before the fluid rotation.

In the following paragraphs, we model the stress dynamics for cells exposed to turbulent cues for the two paradigmatic cases that we employed experimentally: *i*) a continuous solid body rotation (i.e., rolling) with rotation rate Ω = π/*τ*_R_ (no resting phases, *τ*_W_ = 0), and *ii*) multiple, fast reorientations of amplitude π (i.e., flipping) occurring at a rate Ω >> *A*, alternating with resting phases captured by the timescale *τ*_W_ (Fig. 1B).

#### (i) Cell stress generated by rolling

For a population of cells of *H. akashiwo* CCMP452 swimming in a rotating chamber at a constant rotation rate Ω = π/*τ*_R_, where *τ*_R_ is the time taken for half a revolution, a fraction of cells from the population distribution (see **Quantification of the cell stability parameter** above) has a high stability parameter *A* (high mechanical stability) such that these cells swim at a certain angle *θ*_eq_. For this fraction of cells, the continuous rotation does not induce any additional stress above the baseline levels. The remaining cells, which tumble under the continuous rotation, accumulate stress over time at each rotation according to the stress model (Eq. S2). By taking the partial sum in the summation in Eq. S2, the upper envelope of the stress signal over time experienced by the tumbling cells is

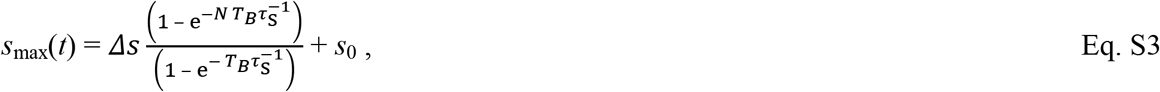

where the sequence of times *t_i_* at which stress is generated is *S* = {*T_B_,…iT_B_,…,NT_B_*} after *N* periodic reorientations. *T_B_* = 2π*B* (*B*^2^Ω^2^ − 1)^−1/2^ is the period of the orbit for the tumbling cells (*S12*), which depends on the stability timescale *B* and on the rotation rate Ω. Note that for *A* = (2*B*)^−1^ << |Ω|, a cell’s trajectory becomes almost circular with period 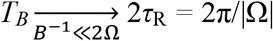 (Fig. S1B). For values of rolling of Ω = 1 rad s^−1^ implemented experimentally in Figs. 3,4 and the stability parameter *A* = 0.09 s of *H. akashiwo* CCMP452 (Fig. S1A), the period of a cell’s trajectory *T_B_* differs from the period of chamber rolling 2π/Ω only by 1%. After multiple reorientations, *N* >>1, the numerator of the first term on the right hand side of Eq. S3 converges to Δ*s*, and the expression is well approximated by

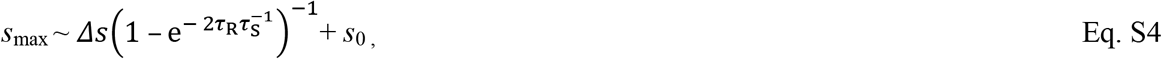

where the cells, by accumulating a relative stress level *s*_max_/*s*_0_ exceeding the stress threshold *h*, switch the direction of migration from upward to downward.

#### (ii) Cell stress generated by flipping

In the case of flipping the chamber at a fast rotation rate Ω_max_ = π rad s^−1^ (*τ*_R_ = 1 s), the condition *A*_max_|Ω_max_| > 1 is satisfied, where *A*_max_ = 1.14 rad s^−1^ is the highest mechanical stability parameter of the population (Fig. 1C). Under these conditions, which corresponds to the case tested experimentally (Fig. 1F), the entirety of the cells in the population would tumble during the flip. Compared to the (slower) continuous rotation case (Fig. 1E), where the emerging downward-migrating is mediated by the distribution of mechanical stabilities in the population, the dynamics for the upward bias present a more pronounced threshold as a function of the resting phase *τ*_W_ because all the cells under this experimental configuration accumulate stress at each flip (Fig. 1F). The accumulation–dissipation dynamics of cellular stress are set by the ratio of the three timescales of the process, the dissipation timescale *τ*_S_ and the period of time between two flips *τ*_W_ + *τ*_R_ (the sum of the resting phase between two flips and the time taken to flip the chamber). We can obtain the maximum stress over time for a cell under multiple periodic reorientations (flips) as

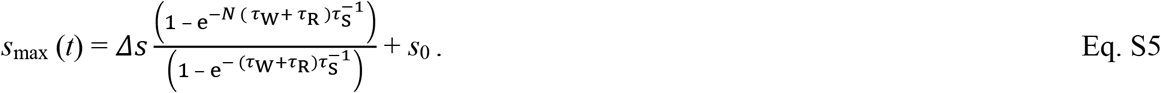

The stress accumulation–dissipation dynamics captured by Eqs. 4 and 5 are plotted in Fig. 3C for the different values of *τ*_W_, which correspond to the range of experimental values for the resting times of Fig. 1F. After many (100) flips, *N* >>1, the numerator of the first term on the right hand side of Eq. S5 converges to Δ*s*, and the expression is well approximated by

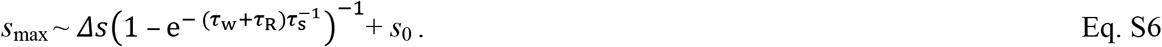

The values of *s*_max_ as a function of the resting time calculated from Eq. S6, which match the experiments presented in Fig. 1F with parameters *N* = 100 flips, a rotation time *τ*_R_ = 1 s, the stress dissipation timescale *τ*_S_ = 87 s from the experiments presented in Fig. 3B, and a value of Δ*s* = 0.5*s*_0_, are plotted in Fig. 3D (black line). Provided that in our experiments *τ*_R_ << *τ*_W_ (except for the case of continuous flipping or rolling with *τ*_W_ = 0 s, which is captured by Eqs. S3 and S4), the expression in Eq. S6 can be further simplified as

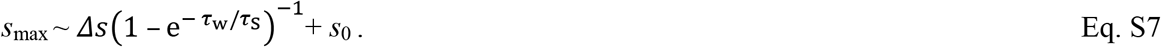

where for τ_S_/τ_W_ >> 1 the first term on the right side becomes much greater than one after rescaling by *s*0, and therefore greater than the stress threshold *h*. In principle, Eq. S2 is applicable to describe the stress dynamics of cells for any turbulent signal (Fig. S2A). The validation of the model on a simplified representation of intermittent turbulence allows its use to investigate the migratory and physiological performance of cells in three-dimensional isotropic turbulent flows (*S13*), for example through direct numerical simulations (*S14, S15*) or turbulence tank experiments (*S16-S18*). Importantly, for a given turbulent signal, the cell’s mechanical stability affects the distribution of the tumbling (Fig. S2B) and resting phases (Fig. S2C), and ultimately determines the migratory behavior of a cell (Fig. 3C). In analogy to the simple case of a gravitactic cell swimming under continuous rolling at a rotation rate Ω considered in Eq. 2 in the main text, where the period *T*B(Ω, *A*) sets the tumbling and therefore the resting time statistics, the characteristic resting time under turbulence is a function *τ*_W_ (*ε, A*) of the turbulent kinetic energy *ε*, which sets the Kolmogorov timescale *τ*_K_ of the microscale turbulent eddies, and of the stability parameter, *A*. To determine the time series of the rotation rate relative to the vertical, Ω(*t*), shown in Fig. 3, we used the time history of the angular orientation of a small passive sphere in homogeneous isotropic turbulence (*S14*), quantified from a direct numerical simulation at Re_λ_ = 65 (time history courtesy of M. Cencini and G. Boffetta), where Re_λ_= u_RMS_ λ/ν is the Taylor Reynolds number, which represents the range of length scales characteristic of a given turbulent flow, with λ = u_RMS_ (15ν/*ε*)^1/2^ the Taylor length scale, u_RMS_ the root-mean-square fluid velocity, and *ε* the turbulent energy dissipation rate.

---

**Fig. S1.**
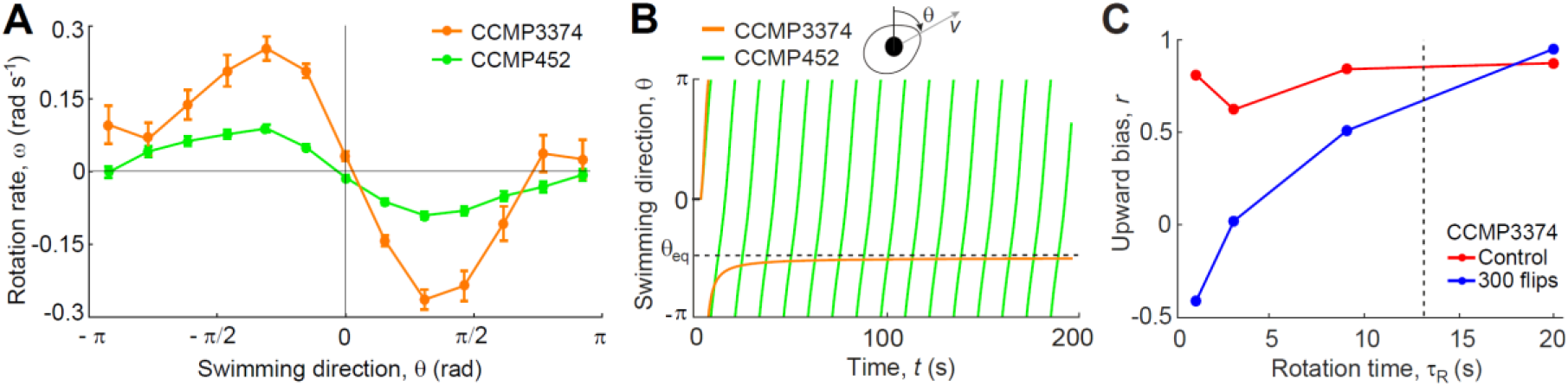
The mechanical stability, together with rotation and resting timescales, determine the magnitude of splitting in CCMP3374. (**A**) Cellular rotation rate, ω, of CCMP452 (green) and CCMP3374 cells (orange), as a function of the cell orientation to the vertical, θ, measured before flipping. Solid lines denote the arithmetic mean over all cell trajectories and error bars represent ± 1 standard error of the mean (s.e.m.). The population stability parameter (CCMP452: A_452_ = 0.09 s^−1^, and CCMP3374: A_3374_ = 0.23 s^−1^) represent the amplitude of the sinusoidal function fitted to the curve (Methods). (**B**) Predicted swimming direction over time for H. akashiwo strains with the two different stability parameters A_452_ and A_3374_ calculated in A, exposed to a solid body rotation with constant angular frequency Ω = 0.2 rad s^−1^. Cells from the CCMP3374 population, with a higher stability parameter, maintain a stable swimming direction (dashed line), θ_eq_ = arcsin(A^−1^Ω), provided (as it is in this case) that the condition A^−1^ |Ω| < 1 is satisfied. Cells from the CCMP452 population tumble and perform periodic orbits (see Eq. 2 in the main text). (**C**) Upward bias, r, in CCMP3374 after N = 300 flips as a function of the rotation time τ_R_ (control cells were kept in quiescent conditions). In agreement with theoretical predictions, the rotation time τ_R_ in combination with the stability parameter A regulate the magnitude of the population split into two subpopulations with opposite mechanical stability (Methods). In particular, we observed that the switch of stability occurred at rotation rates for which A_3374_^−1^|Ω| > 1 (the dashed line satisfies the condition A_3374_^−1^ Ω = 1, with Ω = π/τ_R_).

**Fig. S2.**
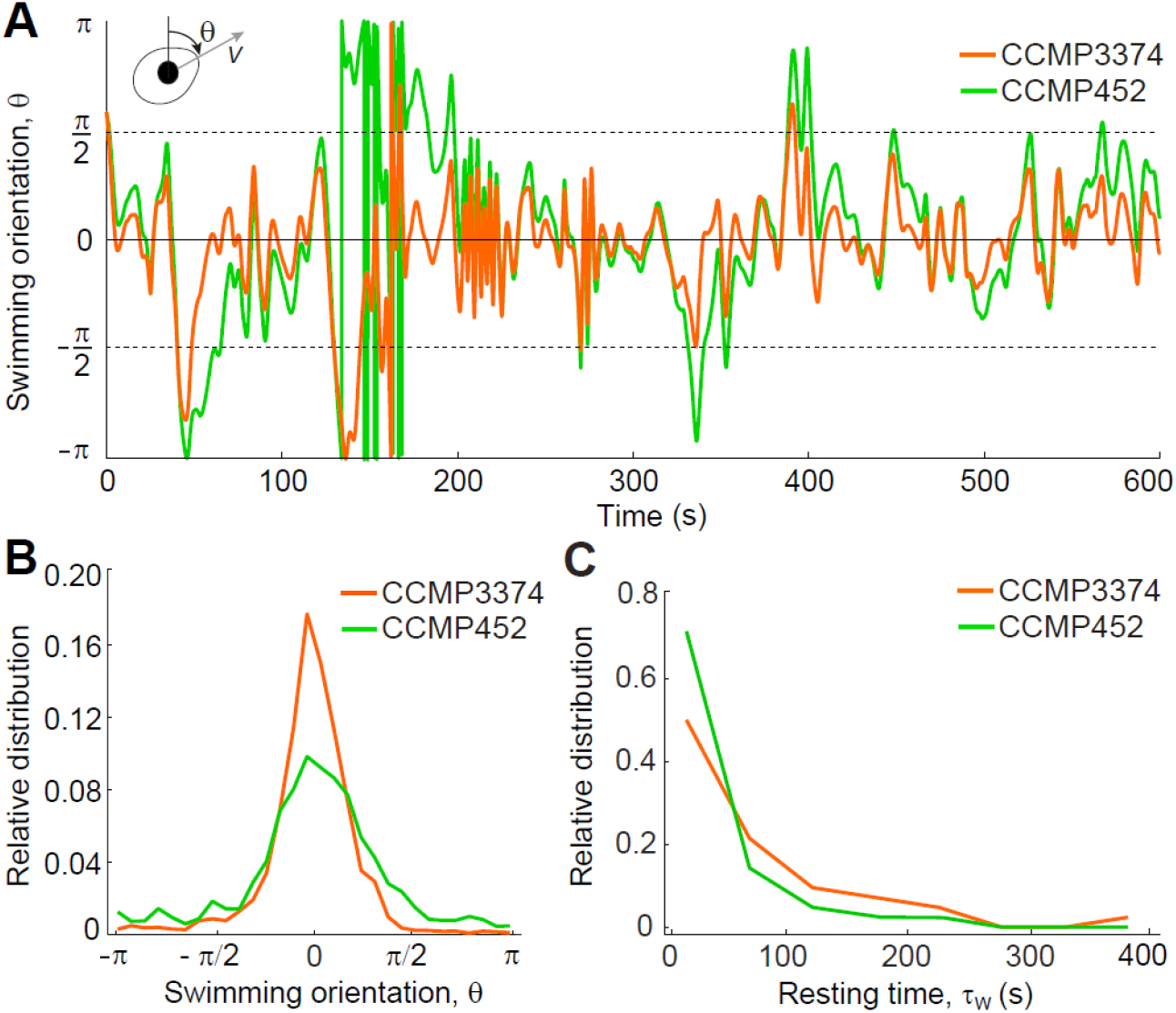
The mechanical stability determines the temporal distribution of tumbling and resting times under turbulence. (**A**) Time series of a cell’s swimming direction relative to gravity, θ(t), calculated from Eq. 1 in the main text, where we imposed the rotation rate Ω(t), which represents the rotation rate relative to the direction of gravity of a passive sphere in a 3D isotropic turbulent flow obtained from a direct numerical simulation (Re_λ_ = 65, ε = 10^−6^ W kg^−1^, ref. S13., Methods). Colored lines are obtained for two values of the stability parameter, A, corresponding to CCMP452 cells (A_452_ = 0.09 s^−1^, green) and CCMP3374 cells (A_3374_ = 0.23 s^−1^, orange). During times when the rotation rate is |Ω|A^−1^< 1, cells are not tumbled but will achieve an equilibrium swimming orientation (Fig. S1B), corresponding in our experiments to a resting time, τ_W_, between reorientations. The cell’s mechanical stability imposes a high-pass filter on the turbulent signal, where CCMP3374 cells, characterized by a higher mechanical stability, regain their equilibrium swimming direction more quickly than CCMP452 cells after being tumbled by turbulence. (**B**) Relative distribution of the swimming direction relative to gravity for CCMP3374 and CCMP452 cells under turbulence. Cells of CCMP3374 characterized by a higher mechanical stability spend longer period of times with an orientation close to their equilibrium orientation in the absence of fluid flow, that is the direction opposite to gravity, θ = 0. (**C**) Relative distribution of the resting time, τ_W_, for CCMP452 cells (green, mean resting time: τ_W452_ = 46 s) and CCMP3374 cells (orange, mean resting time: τ_W3374_ = 104 s). The statistics for the resting times was extracted from the distribution of time periods between two tumbling events, defined as the time points with swimming direction θ = π/2 (dashed lines in A, see Methods).

**Fig. S3.**
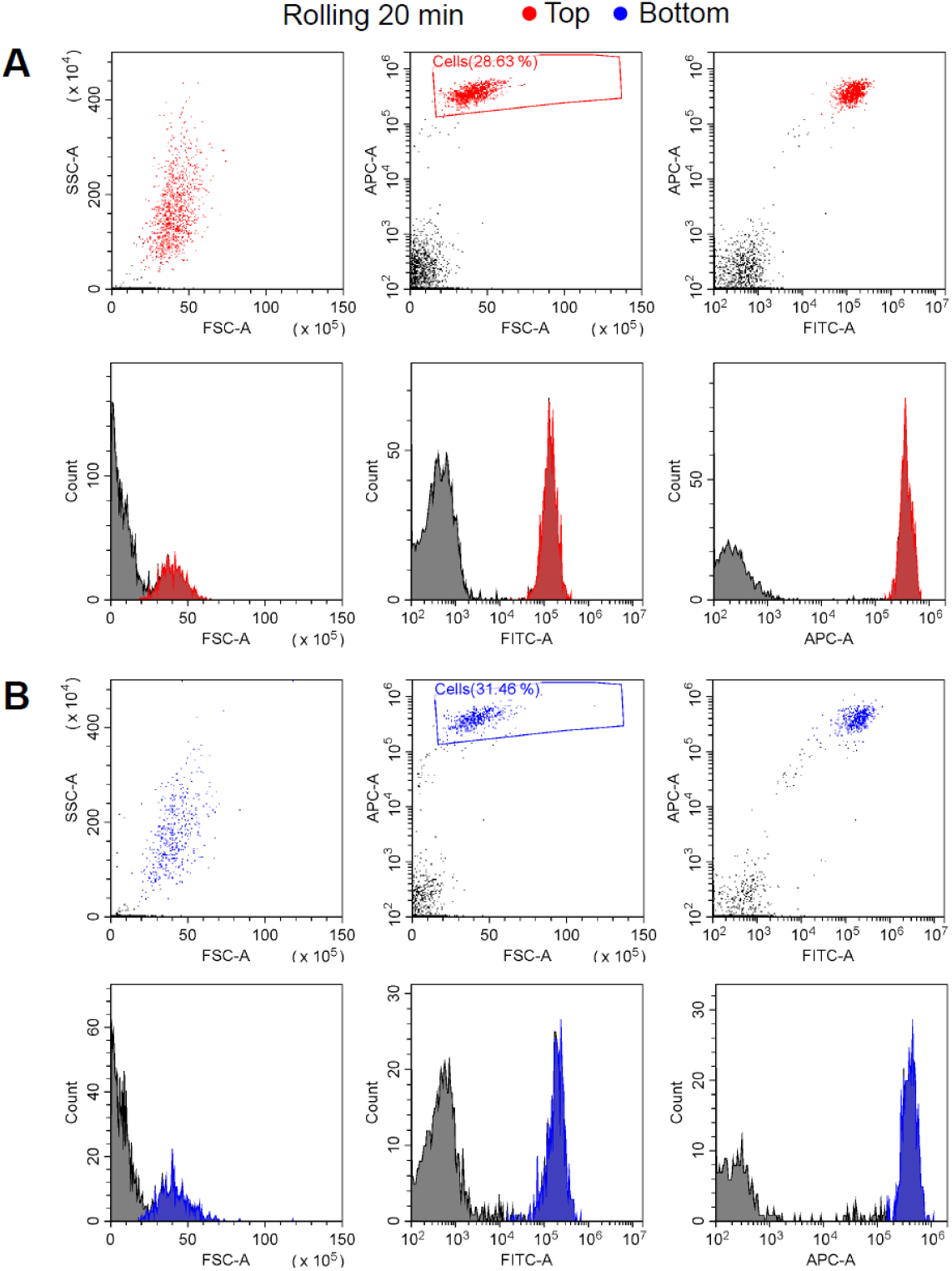
Stress levels in H. akashiwo were higher for the bottom subpopulation. To quantify the accumulation of reactive oxygen species (ROS), cells exposed to rolling in 2 ml cylindrical vials were incubated with 10 μM CM-H_2_DCFDA, a general oxidative stress marker that passively diffuses into live cells and binds to free radicals (see Methods). After a dark-incubation period (30 min), the cells were examined using a flow cytometer. Each sample was run through the flow cytometer until at least 1000 viable cells were detected. The oxidative stress levels are represented as the fluorescence readouts from the flow cytometer in the FITC-A channel, which matches the excitation/emission wavelengths of the CM-H_2_DCFDA dye (Ex/Em: ~488/520 nm). The fluorescence levels were obtained for the turbulence-exposed population extracted from the top (**A**) and from the bottom (**B**) of the cylindrical vials after 20 min rolling. For A and B in the top row, scatter plots show side-scatter (SSC-A) vs. forward-scatter (FSC-A), allophycocyanin (APC-A) vs. FSC-A channels, and APC-A vs. FITC-A channels. The APC-A channel captures the autofluorescence signal from cells in the far-red region of the spectrum (763/43 nm). Each dot within scatterplots represents a single cell. To select only the values in the FITC-A channel from viable H. akashiwo cells for inclusion in the stress analysis (and not bacteria and other debris), we applied a gate using channels FSC-A vs. APC-A to identify cells (red and blue rectangles, % values indicate the relative proportion of cells). In the bottom row, the distributions of cells (colored) and debris (black) are shown for FSC-A, FITC-A, and APC-A. The same staining and flow cytometry protocol was used to quantify stress levels for all the rolling and the H_2_O_2_ experiments.

**Fig. S4.**
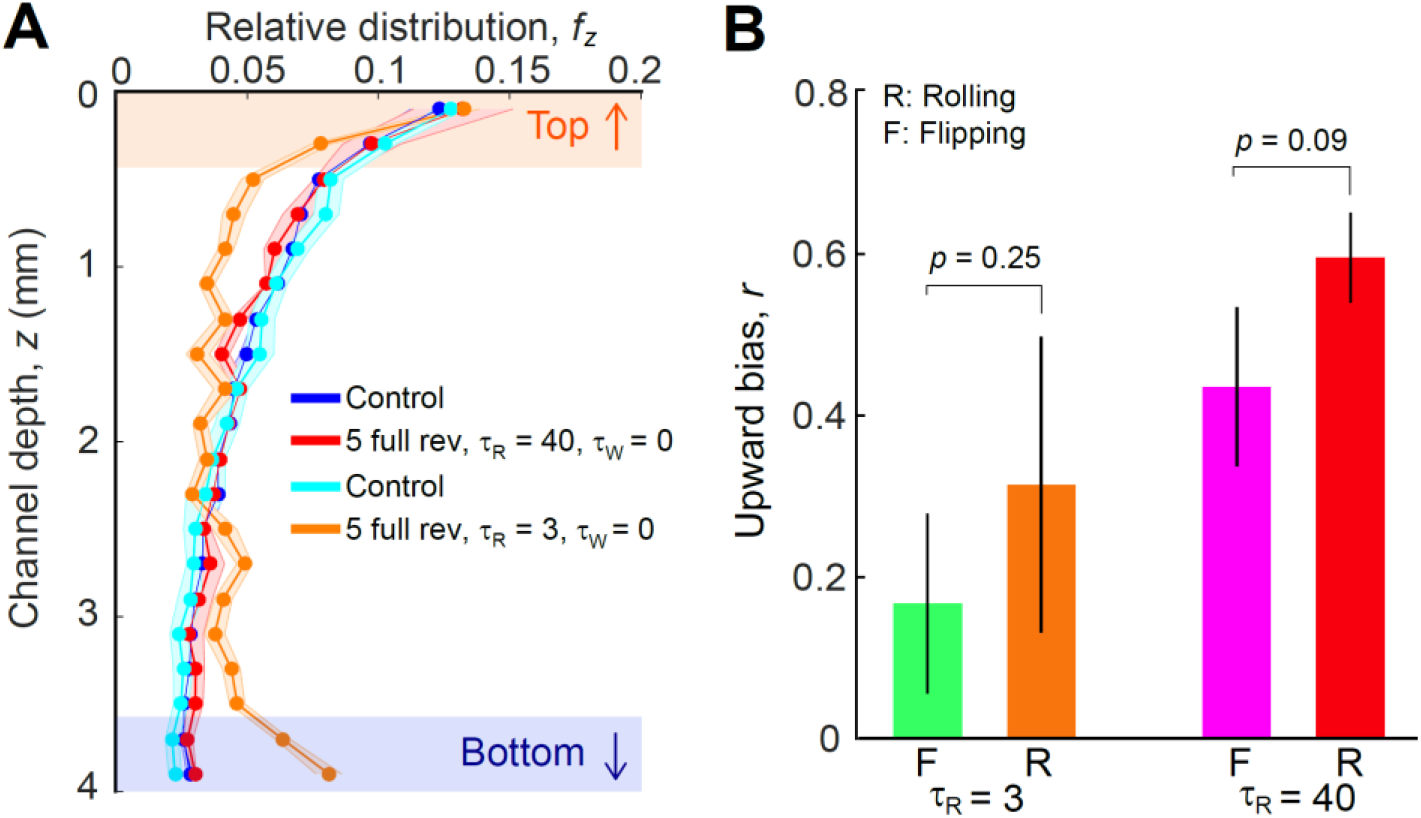
Rolling produced a similar migratory response to that induced by flipping in H. akashiwo. (**A**) The relative distribution of cells in the chamber, f_Z_, was analyzed before (quiescent controls, cyan and blue) and after exposure to continuous rotation (rolling) of the chamber (5 full revolutions with τ_W_ = 0 s) at two different rotation rates (Ω = 1 rad s^−1^, equivalent to a rotation time τ_R_ = 3 s, orange line; Ω = 0.08 rad s^−1^, equivalent to a rotation time τ_R_ = 40 s, red line) following a period of 30 min in which the cell distribution was allowed to equilibrate. Shaded regions in orange and blue represent the top (↑) and the bottom (↓) 400 μm of the chamber, where the concentration of the cells was quantified for the calculation of the upward bias. The upward bias index, r = (f_↑_ − f_↓_)/(f_↑_ + f_↓_), measures the relative proportion of up-swimming (f_↑_) and down-swimming (f_↓_) cells. (**B**) The upward bias obtained in the continuous rolling experiments (R) compared to the flipping experiments (F; N = 10 flips, with the corresponding rotation times and zero resting time, τ_W_ = 0 s). No difference in the upward bias was detected between rolling and flipping (two-sample t-tests; τ_R_ = 3 s rotation time, t_7_ = 1.26, p = 0.25; τ_R_ = 40 s rotation time, t_5_ = 2.09, p = 0.09).

**Fig. S5.**
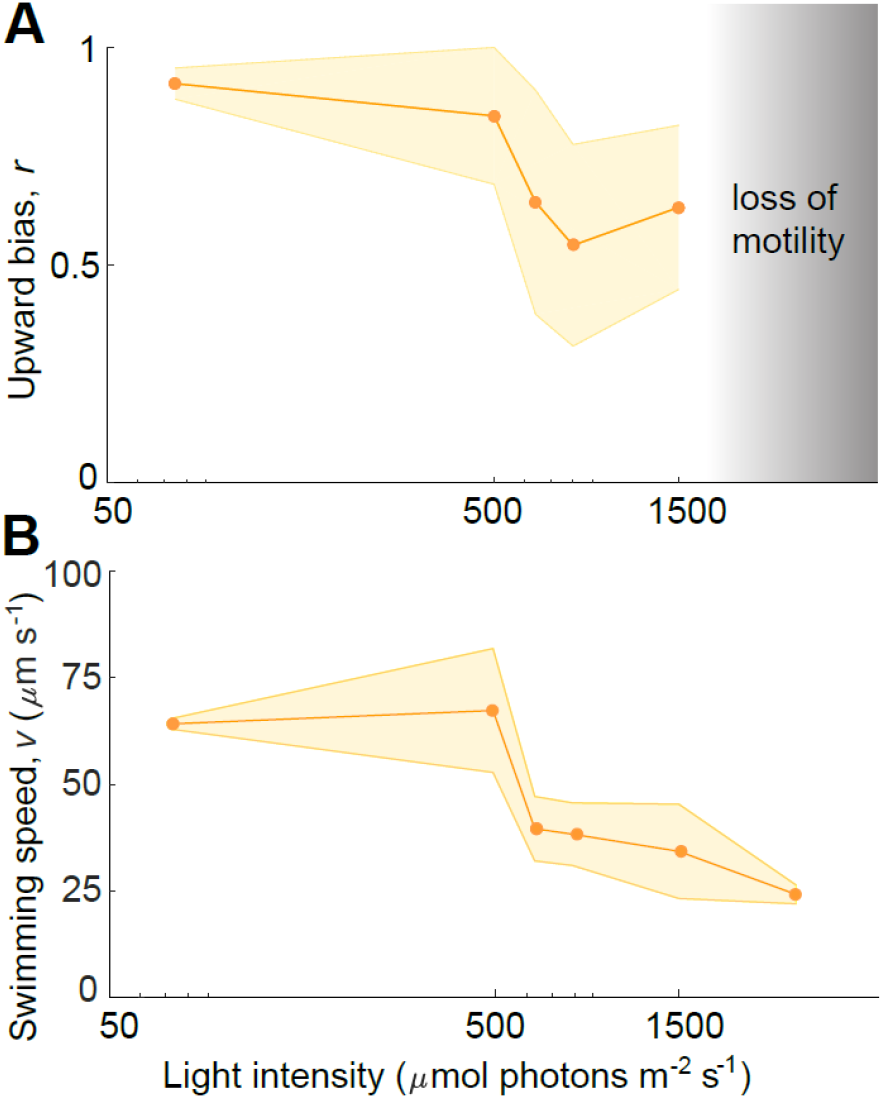
Upward bias (A) and swimming speed (B) of a population of Heterosigma akashiwo as a function of light intensity. We observed a threshold response of the upward bias with increasing light intensity, and a loss of motility (gray shaded region in A) for cells exposed to full spectrum irradiance higher than 650 μmol photons m^−2^ s^−1^. An almost complete loss of motility was observed for light intensities higher than 1500 μmol photons m^−2^ s^−1^, where the values of swimming speed (v = 22 μm s^−1^) were comparable to the sinking speed predicted by the Stokes’ law, v_s_ = 2/9(ρ_p_ − pf)gμ^−1^ R^2^, which for H. akashiwo is v_s_ = 6 μm s^−1^, where (ρ_p_ − pf) = 50 kg m^−3^ is the excess density, g = 9.8 m s^−2^ is the gravitational acceleration, μ = 10^−3^ Pa s is the dynamic viscosity of seawater, and R = 7 μm is the equivalent radius of the cell (S4, S9, S10). The minimum value of light intensity (75 μmol photons m^−2^ s^−1^) corresponds to the intensity used for culturing the population (Methods). Points represent the mean of two replicates and the shaded region is ±1 s.d.

**Fig. S6.**
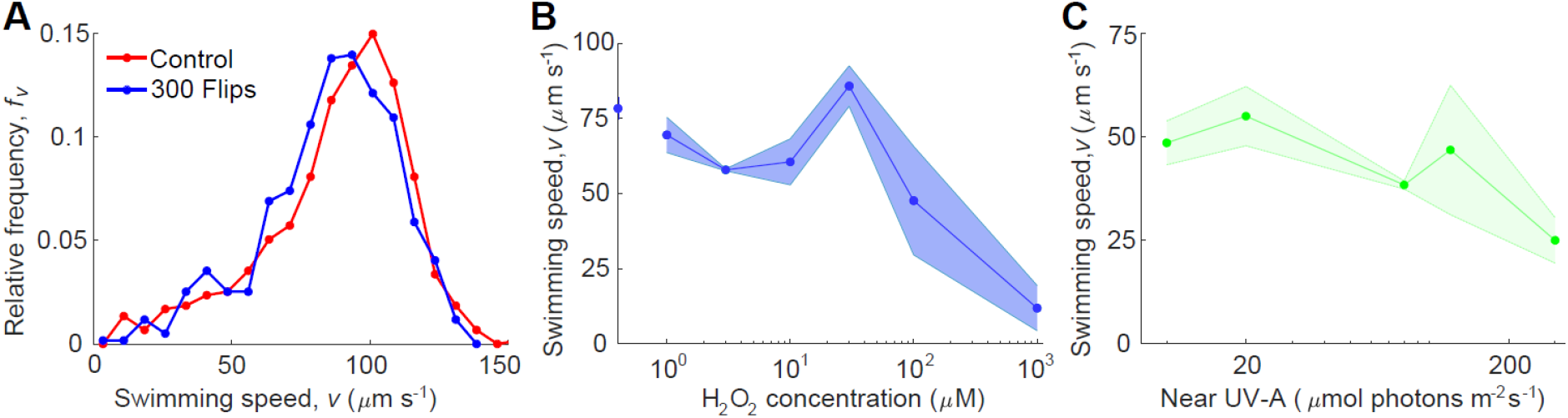
Modulation of swimming speed for H. akashiwo after exposure to different stressors. (**A**) No difference in the distribution of swimming speeds was detected between a population of cells before (control, red curve, v = 88 ± 33 μm s^−1^) and after (blue curve, v = 86 ± 29 μm s^−1^) exposure to N = 300 flips at the highest rotation rate used in our experiments (Ω = 3.14 rad s^−1^, which corresponds to a rotation time τ_R_ = 1 s), with zero resting time between flips (τ_W_ = 0 s). (**B**) Swimming speed after exposure of a population of cells to different concentrations of H_2_O_2_. The blue point on the y-axis indicates the control. A consistent drop in motility was observed at a concentration of 100 μM H_2_O_2_, and a complete loss of motility occurred at a concentration of 1 mM H_2_O_2_, where the detected swimming speed, v = 11 μm s^−1^, was comparable to the sinking speed predicted by the Stokes’ law, v_s_ = 2/9(ρ_p_ − ρfgμ^−1^R^−2^, which for H. akashiwo is v_s_ = 6 μm s^−1^, where (ρ_p_ − ρ_f_ = 50 kg m^−3^ is the excess density, g = 9.8 m s^−2^ is the gravitational acceleration, μ = 10^−3^ Pa s is the dynamic viscosity of seawater, and R = 7 μm is the equivalent radius of the cell (S4, S9, S10). Points represent the mean of three replicates and the shaded region is ± 1 s.d. (**C**) Swimming speed after exposure of a population of cells to different UV-A intensities. Cells maintained normal motility after exposure up to an intensity of 120 μmol photons m^−2^ s^−1^ UV-A. A drop in motility was detected at a photon flux density of 300 μmol photons m^−2^ s^−1^ UV-A. Points represent the mean of three replicates and the shaded region is ± 1 s.d.

**Table S1.**
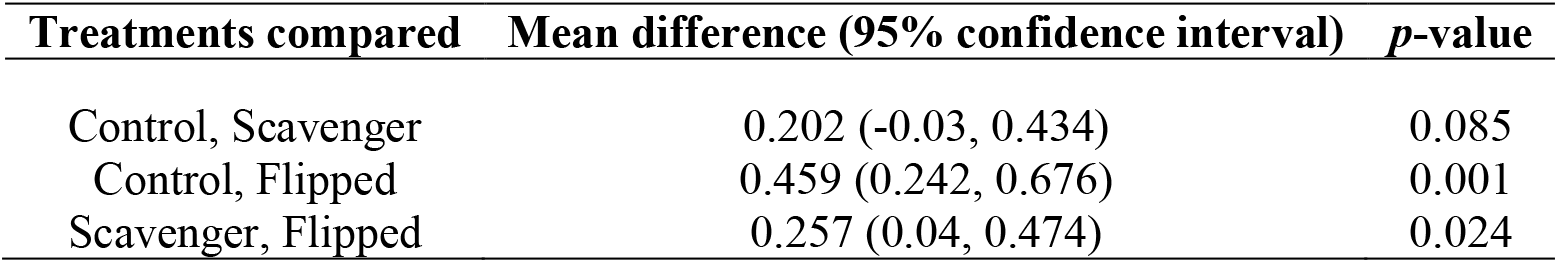
Multiple comparisons analysis (Tukey’s HSD test) of the upward bias of *H. akashiwo* CCMP452 among a population that was flipped, a population that was flipped after having been cultured in the presence of a scavenger of reactive oxygen species (potassium iodide), and a control population (without flipping and no potassium iodide). Control = quiescent control; Scavenger = a population grown in f/2 medium with potassium iodide at a concentration of 100 μM, flipped 100 times *τ*_R_ = 3 s*, τ*_W_ = 0 s; Flipped = a population grown in f/2 medium, flipped 100 times (*τ*_R_ = 3 s, *τ*_W_ = 0 s).

**Table S2.**
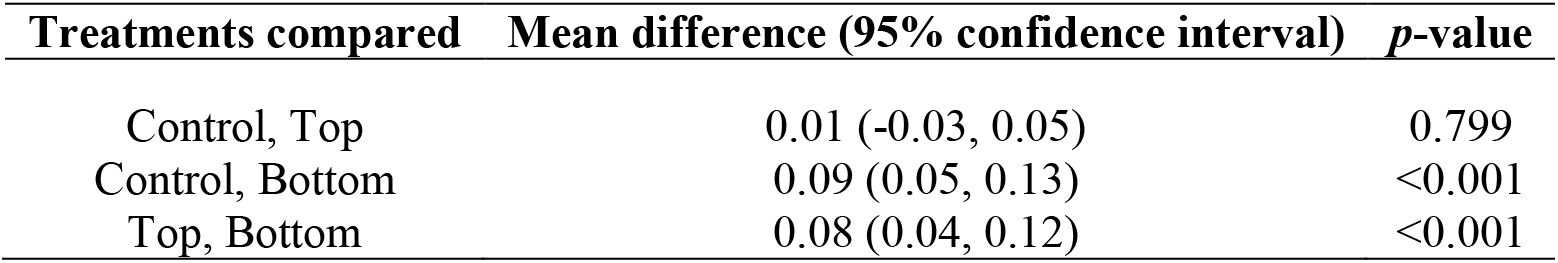
Multiple comparisons analysis (Tukey’s HSD test) of the photosynthetic quantum yield *H. akashiwo* CCMP452 among subpopulations that were collected from the top and bottom after rolling, and a control population (without rolling). Control = quiescent control; Top = subpopulation harvested from the top of the chamber, 5 min rolling time (Ω = 1 rad s^−1^); Bottom = subpopulation harvested from the bottom of the chamber, 5 min rolling time (Ω = 1 rad s^−1^).

| Treatments compared | Mean difference (95% confidence interval) | *p*-value |
| --- | --- | --- |
| Control, Top | 0.01 (-0.03, 0.05) | 0.799 |
| Control, Bottom | 0.09 (0.05, 0.13) | <0.001 |
| Top, Bottom | 0.08 (0.04, 0.12) | <0.001 |

## References

1. M. M. Omand, E. A. D’Asaro, C. M. Lee, M. J. Perry, N. Briggs, I. Cetinić, A. Mahadevan, Eddy-driven subduction exports particulate organic carbon from the spring bloom, Science 348, 222–225 (2015).

2. H. Alexander, M. Ruoco, S. T. Haley, S. T. Wilson, D. M. Karl, S. T. Dyhrman, Functional group-specific traits drive phytoplantkon dyanmics in the oligotrophic ocean, Proc. Natl. Acad. Sci. U.S.A. 112, E5972–E5979 (2015).

3. F. d’Ovidio, S. De Monte, S. Alvain, Y. Dandonneau, M. Lévy, Fluid dynamical niches of phytoplankton types, Proc. Natl. Acad. Sci. U.S.A. 107, 18366–18370 (2010).

4. R. Margalef, Turbulence and marine life, Sci. Mar. 61, 109–123 (1997).

5. A. D. Barton, B. A. Ward, R. G. Williams, M. J. Follows, The impact of fine-scale turbulence on phytoplankton community structure, Fluid. Environ. 4, 34–49 (2014).

6. L. Karp-Boss, E. Boss, P. A. Jumars, Nutrient fluxes to planktonic osmotrophs in the presence of fluid motion, Oceanogr. Mar. Biol. 34, 71–107 (1996).

7. J. S. Guasto, R. Rusconi, R. Stocker, Fluid Mechanics of planktonic microorganisms, Ann. Rev. Fluid Mech. 44, 373–400 (2012).

8. F. Peters, C. Marrasé, Effects of turbulence on plankton: an overview of experimental evidence and some theoretical considerations, Mar. Ecol. Prog. Ser. 205, 291–306 (2000).

9. M. J. Zirbel, F. Veron, M. I. Latz, The reversible effect of flow on the morphology of *Ceratocorys horrida* (Peridiniales, Dinophyta), J. Phycol. 36, 46–58 (2000).

10. E. Berdalet, F. Peters, V. L. Koumandou, C. Roldan, O. Gudayol, M. Estrada, Species–specific physiological response of dinoflagellates to quantified small-scale turbulence, J. Phycol. 43, 965–977 (2007).

11. W. M. Durham, E. Climent, M. Barry, F. De Lillo, G. Boffetta, M. Cencini, R. Stocker, Turbulence drives microscale patches of motile phytoplankton, Nat. Commun. 4, 2148 (2013).

12. R. E. Brier, C. C. Lalescu, D. Waas, M. Wilczek, M. G. Mazza, Emergence of phytoplankton patchiness at small scales in mild turbulence, Proc. Natl. Acad. Sci. U.S.A. 115, 12112–12117 (2018).

13. A. Amato, G. Dell’Aquila, F. Musacchia, R. Annunziata, A. Ugarte, N. Maillet, A. Carbone, M. Ribera d’Alcalá, R. Sanges, D. Iuducone, M. I. Ferrante, Marine diatoms change their gene expression profile when exposed to microscale turbulence under nutrient replete conditions, Sci. Rep. 7, 3826 (2017).

14. A. Falciatore, M. Ribera d’Alcalá, P. Croot, C. Bowler, Perception of environmental signals by a marine diatom, Science 288, 2363–2366 (2000).

15. B. J. Gemmell, G. Oh, E. J. Buskey, T. A. Villareal, Dynamic sinking behavior in marine phytoplankton: rapid changes in buoyancy may aid in nutrient uptake, Proc. R. Soc. B 283, 20161126 (2016).

16. A. Sengupta, F. Carrara, R. Stocker, Phytoplankton can actively diversify their migration strategy in response to turbulence cues, Nature 543, 555–558 (2017).

17. T. A. Villareal, E. J. Carpenter, Buoyancy regulation and the potential for vertical migration in the oceanic cyanobacterium *Trichodesmium*, Microb. Ecol. 45, 1–10 (2003).

18. J. M. Sullivan, P. L. Donaghay, J. E. B. Rines, Coastal thin layers dynamics: consequences to biology and optics, Cont. Shelf Res. 30, 50–65 (2010).

19. P. A. Jumars, J. H. Trowbridge, E. Boss, L. Karp-Boss, Turbulence-plankton interaction: a new cartoon, Mar. Ecol. 30, 133–150 (2009).

20. R. Watteaux, G. Sardina, L. Brandt, D. Iudicone, On the time scales and structure of Lagrangian intermittency in homogeneous isotropic turbulence, J. Fluid Mech., 867, 438–481 (2019).

21. T. J. Pedley, J. O. Kessler, Hydrodynamic phenomena in suspensions of swimming microorganisms, Ann. Rev. Fluid Mech. 24, 313–358 (1992).

22. Y. Goulev, S. Morlot, A. Matisfas, B. Huang, M. Molin, M. B. Toledano, G. Charvin, Nonlinear feedback drives homeostatic plasticity in H_2_O_2_ stress response, eLife 23971 (2017).

23. A. Vardi, D. Eisenstadt, O. Murik, I. Berman-Frank, T. Zohary, A. Levine, A. Kaplan, Synchronization of cell death in a dinoflagellate population is mediated by an excreted thiol protease, Env. Microb. 9, 360–369 (2006).

24. S. G. van Creveld, S. Rosenwasser, D. Schatz, I. Koren, A. Vardi, Early perturbation in mitochondria redox homeostasis in response to environmental stress predicts cell fate in diatoms, ISME 9, 385–395 (2015).

25. B. D’Autréaux, M. B. Toledano, ROS as signaling molecules: mechanisms that generate specificity in ROS homeostasis, Nat. Rev. Mol. Cell Biol. 8, 813–824 (2007).

26. J. W. Rijstenbil, Assessment of oxidative stress in the planktonic diatom *Thalassiosira pseudonana* in response to UVA and UVB radiation, J. Plank. Res. 24, 1277–1288 (2002).

27. D. J. Mcgillicuddy Jr., D. M. Anderson, D. R. Lynch, D. W. Townsend, Mechanisms regulating large-scale seasonal fluctuations in *Alexandrium fundyense* in the Gulf of Maine: Results from physical-biological model, Deep-Sea Res. II 52, 2698–2714 (2005).

28. A. Mizrachi, S. Graff van Creveld, O. H. Shapiro, S. Rosenwasser, A. Vardi, Light-dependent single-cell heterogeneity in the chloroplast redox state regulates cell fate in a marine diatom, eLife 47732 (2019).

29. J. Waring, M. Klenell, U. Bechtold, G. J. C. Underwood, N. R. Baker, Light-induced responses of oxygen photoreduction, reactive oxygen species production and scavenging in two diatom species, J. Phycol. 46, 1206–1217 (2010).

30. Y. Nishiyama, H. Yamamoto, S. I. Allakhverdiev, M. Inaba, A. Yokota, N. Murata, Oxidative stress inhibits the repair of photodamage to the photosynthetic machinery, EMBO J. 20, 5587–5594 (2001).

31. S. Rosenwasser, S. Graff van Creveld, D. Schatz, S. Malitsky, O. Tzfadia, A. Aharoni, Y. Levin, A. Gabashvili, E. Feldmesser, A. Vardi, Mapping the diatom redox-sensitive proteome provides insight into response to nitrogen stress in the marine environment, Proc. Natl. Acad. Sci. U.S.A. 111, 2740–2745 (2014).

32. N. R. Baker, Chlorophyll fluorescence: a probe of photosynthesis in vivo, Ann. Rev. Plant Biol. 59, 89–113 (2008).

33. J. Lämke, I. Bäurle, Epigenetic and chromatin-based mechanisms in environmental stress adaptation and stress memory in plants, Gen. Biol. 18, 124 (2017).

34. S. Colabrese, K. Gustavsson, A. Celani, L. Biferale, Flow navigation by smart microswimmers via reinforcement learning, Phys. Rev. Lett. 118, 158004 (2017).

35. M. K. Thomas, C.T. Kremer, C.A. Klausmeier, E. Lichtman, A global pattern of thermal adaptation in marine phytoplankton, Science, 338, 1085–1088 (2012).

36. A. D. Barton, A. J. Irwin, Z. V. Finkel, C. A. Stock, Anthropogenic climate change drives shift and shuffle in North Atlantic phytoplankton communities, Proc. Natl. Acad. Sci. U.S.A. 113, 2964–2969 (2016).

37. C. J. Gobler, O. M. Doherty, T. K. Hattenrath-Lehmann, A. W. Griffith, Y. Kang, R. W. Litaker, Ocean warming since 1982 has expanded the niche of toxic algal blooms in the North Atlantic and North Pacific oceans, Proc. Natl. Acad. Sci. U.S.A. 114, 4975–4980 (2017).

38. Y. Hara, M. Chihara, Morphology, ultrastructure and taxonomy of the raphidophycean alga *Heterosigma akashiwo*, Bot. Mag. 100, 151–163 (1987).

39. M. Wada, A. Miyazaki, T. Fujii, On the mechanisms of diurnal vertical migration behavior of *Heterosigma akashiwo* (Raphidophyceae), Plant Cell. Physiol. 26, 431–436 (1985).

40. U. Sheyn, S. Rosenwasser, S. Ben-Dor, Z. Porat, A. Vardi, Modulation of host ROS metabolism is essential for viral infection of a bloom-forming coccolithophore in the ocean, ISME J. 10, 1742–54 (2016).

41. C. Dunand, M. Crèvecoeur, C. Penel, Distribution of superoxide and hydrogen peroxide in *Arabidopsis* root and their influence on root development: possible interaction with peroxidases. New Phyt. 174, 332–341 (2007).

42. R. Martínez, E. Orive, A. Laza-Martínez, S. Seoane, Growth response of six strains of *Heterosigma akashiwo* to varying temperature, salinity and irradiance conditions. J. Plank. Res. 32, 529–538 (2010).

43. E. Trampe, J. Kolbowski, U. Schreiber, M. Kühl, Rapid assessment of different oxygenic phototrophs and single-cell photosynthesis with multicolour variable chlorophyll fluorescence imaging, Mar. Biol. 158, 1667–1675 (2011).

44. W. Bilger, O. Bjoerkman, Role of the xanthophyll cycle in photoprotection elucidated by measurements of light-induced absorbance changes, fluorescence and photosynthesis in leaves of *Hedera canariensis*, Photosynth. Res. 25, 173–185 (1990).

45. A. M. Roberts, Geotaxis in motile microorganisms, J. Exp. Biol. 53, 687–699 (1970).

46. C. L. Dupont, T. J. Goepfert, P. Lo, L. Wei, B. A. Ahner, Diurnal cycling of glutathione in marine phytoplankton: Field and culture studies, Limnol. Oceanogr. 49, 991–996 (2004).

## Supplementary References

S1. S. A. Thorpe, An introduction to ocean turbulence (Cambridge Univ. Press, Cambridge, MA, 2007).

S2. H. L. Fuchs, G. P. Gerbi, Seascape-level variation in turbulence- and wave-generated hydrodynamic signals experienced by phytoplankton, Prog. Ocean. 141, 109–129 (2016).

S3. K. W. Foster, R. D. Smyth, Light Antennas in phototactic algae, Microbiol. Rev. 44, 572–630 (1980).

S4. A. Sengupta, F. Carrara, R. Stocker, Phytoplankton can actively diversify their migration strategy in response to turbulence cues, Nature 543, 555–558 (2017).

S5. N. Oulette, H. Xu, E. Bodenschatz, A quantitative study of three-dimensional Lagrangian particle tracking algorithms, Exp. Fluids 40, 301–313 (2006).

S6. T. Fenchel, How dinoflagellates swim, Protist 152, 329–338 (2002).

S7. A. M. Roberts, Geotaxis in motile microorganisms, J. Exp. Biol. 53, 687–699 (1970).

S8. H. M. Shapiro, Practical Flow Cytometry, (Wiley & Sons, New York, NY, 2003).

S9. Y. Hara, M. Chihara, Morphology, ultrastructure and taxonomy of the raphidophycean alga *Heterosigma akashiwo*, Bot. Mag. 100, 151–163 (1987).

S10. M. Wada, A. Miyazaki, T. Fujii, On the mechanisms of diurnal vertical migration behavior of *Heterosigma akashiwo* (Raphidophyceae), Plant Cell. Physiol. 26, 431–436 (1985).

S11. K. Mukai, Y. Shimasaki, X. Qiu, Y. Kato-Unoki, K. Chen, M. R. M. Khanam, Y. Ohima, Effects of light and hydrogen peroxide on gene expression of newly identified antioxidant enzymes in the harmful algal bloom species *Chattonella marina*, Eur. J. Phyc. 54, 393–403 (2019).

S12. T. J. Pedley, J. O. Kessler, Hydrodynamic phenomena in suspensions of swimming microorganisms, Ann. Rev. Fluid Mech. 24, 313–358 (1992).

S13. S. B. Pope, Turbulent flows (Cambridge Univ. Press, Cambridge, MA, 2000)

S14. W. M. Durham, E. Climent, M. Barry, F. De Lillo, G. Boffetta, M. Cencini, R. Stocker, Turbulence drives microscale patches of motile phytoplankton, Nat. Commun. 4, 2148 (2013).

S15. R. E. Brier, C. C. Lalescu, D. Waas, M. Wilczek, M. G. Mazza, Emergence of phytoplankton patchiness at small scales in mild turbulence, Proc. Natl. Acad. Sci. U.S.A. 115, 12112–12117 (2018).

S16. M. J. Zirbel, F. Veron, M. I. Latz, The reversible effect of flow on the morphology of *Ceratocorys horrida* (Peridiniales, Dinophyta), J. Phycol. 36, 46–58 (2000).

S17. E. Berdalet, F. Peters, V. L. Koumandou, C. Roldan, O. Gudayol, M. Estrada, Speciesspecific physiological response of dinoflagellates to quantified small-scale turbulence, J. Phycol. 43, 965–977 (2007).

S18. A. Amato, G. Dell’Aquila, F. Musacchia, R. Annunziata, A. Ugarte, N. Maillet, A. Carbone, M. Ribera d’Alcala, R. Sanges, D. Iuducone, M. I. Ferrante, Marine diatoms change their gene expression profile when exposed to microscale turbulence under nutrient replete conditions, Sci. Rep. 7, 3826 (2017).

